# Benchmarking PWM and SVM-based Models for Transcription Factor Binding Site Prediction: A Comparative Analysis on Synthetic and Biological Data

**DOI:** 10.1101/2025.03.20.644354

**Authors:** Manuel Tognon, Alisa Kumbara, Andrea Betti, Lorenzo Ruggeri, Rosalba Giugno

**Affiliations:** Computer Science Department, University of Verona, Verona, Italy; Department of Engineering for Innovation Medicine, University of Verona, Verona Italy; Department of Computer, Control, and Management Engineering “Antonio Ruberti”, University of Rome “La Sapienza”, Rome Italy

## Abstract

Transcription Factors (TFs) are essential regulatory proteins that control the cellular transcriptional states by binding to specific DNA sequences known as Transcription Factor Binding Sites (TFBSs) or motifs. Accurate TFBS identification is crucial for unraveling regulatory mechanisms driving cellular dynamics. Over the years, various computational approaches have been developed to model TFBSs, with Position Weight Matrices (PWMs) being one of the most widely adopted methods. PWMs provide a probabilistic framework by representing nucleotide frequencies at every position within the binding site. While effective and interpretable, PWMs face significant limitations, such as their inability to capture positional dependencies or model complex interactions. To address these, advanced methods, such as Support Vector Machine (SVM)-based models, have been introduced. Leveraging human ChIP-seq data from ENCODE, this study systematically benchmarks the predictive performance of PWM and SVM-based models across different scenarios. We evaluate the impact of key factors such as training dataset size, sequence length, and kernel functions (for SVMs) on models’ performance. Additionally, we explore the impact of synthetic versus real biological background data during model training. Our analysis highlights strengths and limitations of both PWM and SVM-based approaches under different conditions, providing practical guidance for selecting and tailoring models to specific biological datasets. To complement our analysis, we present a comprehensive database of pretrained SVM models for TFBS detection, trained on human ChIP-seq data from diverse cell lines and tissues. This resource aims to facilitate broader adoption of SVM-based methods in TFBS prediction and enhance their practical utility in regulatory genomics research.

## 1 Introduction

Transcription factors (TFs) are key regulatory proteins that orchestrate different cellular processes, including transcriptional regulation, cellular differentiation, and development [1—3]. TFs execute their regulatory function by binding to short DNA sequences called transcription factor binding sites (TFBSs). TFBS, often also referred to as *motifs*, are distributed across different genomic regions, encompassing promoters, enhancers, silencers, or insulators, and even within coding sequences [4—7]. The precise configuration of TFBS sequences govern TFs’ regulatory function [8,9], highlighting the significance and importance of identifying putative binding sites for elucidating the intricate regulatory mechanisms underlying the cellular dynamics.

Different experimental methods have been developed to identify TFBS both in living cells (*in vivo*) and controlled environments using purified components [10]. Earlier assays, such as electrophoretic mobility shift assay [11] and footprinting [12] focused on small datasets of target sequences, while high-throughput techniques like PBM, SELEX, and ChIP [13—16] enabled broader analyses.

ChIP assays [15,16] transformed TFBS research, allowing genome-wide identification of bound regions *in vivo*. In ChIP, TF–DNA complexes are cross-linked with formaldehyde, fragmented, and immunoprecipitated using TF-specific antibodies. Following cross-link reversal, the DNA fragments are amplified and analyzed via microarray hybridization (ChIP-on-Chip [15]) or sequencing (ChIP-seq [16,17]). Fragments are then mapped to the genome, where peak-calling algorithms [18—20] identify binding regions by detecting significant enrichment compared to control samples. ChIP-seq produces large datasets, often covering tens of thousands of genomic regions, each ranging from hundreds to thousands of nucleotides. This data enables the inference of potential TFBS locations, establishing ChIP-seq as the current *in vivo* “gold standard” method. However, it has limitations: ChIP-seq may detect indirect binding from other TFs, include false positives due to antibody quality, and yield low-resolution data with excess flanking nucleotides [21—23]. ChIP-exo [24] addresses this by using a lambda exonuclease to trim excess flanking nucleotides. Since most TFs bind to target sequences in open chromatin regions, methods like DNase-seq [25] can be used to identify potential TFBS *in vivo*, particularly when the target TF is unknown.

Databases like ENCODE, Cistrome, and GTRD [26—28] offer extensive collections of ChIP-seq datasets across various organisms, tissues, and cell types. ENCODE [26] provides a collection of ChIP-seq and DNase-seq data on human functional elements from different tissues and cell types.

Despite these resources, identifying precise binding sites within sequences remains challenging [29,30]. Motif discovery algorithms [23,31,32] help bridge this gap by analyzing sequence datasets to learn TF binding affinities and identify potential TFBS locations. The learned binding affinities are then encoded in computational models representing the discovered TFBSs. Position Weighted Matrices (PWMs) [33,34] are robust and interpretable models quantifying the contribution of each nucleotide at each motif position. Despite their effectiveness and widespread adoption, PWMs have some limitations. They assume positional independence within TFBS motifs, accommodating different binding site configurations [35], but potentially increasing susceptibility to noise, which can result in false positive predictions of binding sites [23]. Alternative models have been proposed, aiming to address these limitations.

Support Vector Machine (SVMs) [36] have been employed to address different challenges in bioinformatics [37], including motif discovery and modeling. SVM-based motif models [23,38] leverage *k*-mers frequencies within input sequences to create a classification space, distinguishing bound and unbound sequences. While studies demonstrated that SVM-based models can outperform PWMs in predictive tasks [39,40], they come with limitations. SVM-based models are limited to capturing short *k*-mers (10-12 bp), and their performance heavily relies on training data quality [23].

Deep neural networks (DNNs) gained prominence in computational biology [41,42] due to their ability to capture complex patterns. Convolutional neural networks (CNNs) have been applied to model TF-DNA interactions. By leveraging nonlinear transformations, CNNs learn complex sequence features underlying TFBSs. Tools such as DeepBind [43] or Basset [44] employed CNNs tailored for motif discovery.

However, DNNs require large training datasets and substantial computational resources The lack of interpretability of their models [45], makes difficult deriving biological insights. In contrast, PWM and SVM-based models offer scalable, hardware-independent and more interpretable solutions.

Motif discovery algorithms face challenges when applied to large datasets, including scalability issues and noise susceptibility. To mitigate these issues, motif discovery tools are run on subsets of the input sequences [46,47]. While this technique can reduce computational demands, it may produce inaccurate or biased models [48]. Another technique to minimize noise in ChIP-seq sequences involves restricting the width of the input sequences, focusing on regions surrounding the peak summit [49]. However, this method may overlook TFBS occurrences distal from the peak summit. For methods employing SVMs, an additional factor is the selection of the kernel function, as it determines the model’s ability to learn the patterns defining TFBSs [50].

Several studies have evaluated the performance of TFBS models [51—54], but many rely on using synthetic datasets [51] or simpler genomes [52] in their evaluations, limiting their applicability to complex regulatory landscapes.

In this paper, we compare the predictive performance of PWM and SVM-based motif models (**Section 2** and **3**) trained using three widely used motif discovery tools: MEME [55—57], STREME [58], and LS-GKM [59]. Models were trained on human ChIP-seq data from ENCODE [26], focusing on TFs representing all DNA-binding domains (DBDs) [1] (**Section 4**). We evaluate the impact of training dataset size and sequence width, kernel functions and the impact of background data on models’ accuracy, comparing synthetic (shuffled input sequences) and biologically feasible (DNase-seq regions) backgrounds (**Section 5**). We further explore the impact of dataset imbalance on models’ performance, simulating a ‘needle in the haystack’ scenario [60].

While SVM-based motif models potentially offer notable advantages, their adoption has been hindered by the lack of accessible pretrained models, unlike PWMs, which benefit from established resources. To address this gap, we present a comprehensive database of SVM-based motif models trained on human ChIP-seq data obtained on diverse cell lines and tissues from ENCODE (**Section 6**).

Through these comparisons, we deliver valuable guidance for selecting the most suitable motif model based on data characteristics and research goals, while introducing a resource enhancing the accessibility and utility of SVM-based models.

## 2 Position Weight Matrix Motif Models

Position Weight Matrices (PWMs) [33,34] are widely recognized as the predominant models for encoding TFs’ binding preferences. PWMs provide an additive framework describing the contribution of each binding site position to TF-DNA binding events. PWMs achieve this by tallying the frequency of each nucleotide at every TFBS position, derived computing an ungapped alignment profile constructed using the sequences identified as potential motifs by motif discovery algorithms [23]. PWMs are statistically evaluated using metrics, such as nucleotide conservation or information content [61]. PWMs are often visualized as sequence logos [62], where the height of each nucleotide corresponds to its information content., PWMs have become the standard model for representing and visualizing TFBS motifs, with databases like JASPAR [63] or HOCOMOCO [64], providing annotated PWMs covering a broad spectrum of known binding sites. However, PWMs come with limitations. PWMs assume positional independence within the motif, potentially causing the identification of false negative or false positive TFBSs [23,35]. Despite this, several algorithms have been proposed to discover TFBS in genomic sequences, constructing PWMs to represent the identified motifs. Methods utilizing enumeration or alignment of motif candidate sequences have gained attention due to their balance between computational efficiency and model interpretability. While more complex methods, such as those leveraging Deep Neural Networks (DNNs) [42] can yield PWMs, they come with certain trade-offs between computational requirements for training and model interpretability. On the other hand, methods enumerative and alignment-based methods maintain interpretability while offering a more computationally scalable framework [65].

### 2.1 Enumerative Motif Discovery for PWM Training: An Example with STREME

The recently introduced algorithm STREME [58] employs an enumerative-based approach to discover TFBS motifs in DNA sequences (**Supplementary File Section 1**). Enumeration-based motif discovery methods search for overrepresented sequence patterns within the input sequence dataset, relative to a background set of sequences [23]. By counting the approximate frequency of each sequence fragment in the foreground (positive) and background (negative) datasets, these methods evaluate whether the observed differences in occurrence are statistically significant, enabling the discovery of enriched motifs. Enumerating subsequences and comparing their frequencies can be computationally demanding. Therefore, many enumerative algorithms employ optimized data structures like suffix trees (STs), to enhance this process. STREME leverages STs to enhance computational efficiency and perform approximate sequence matching to discover enriched patterns. By constructing a ST from positive sequences, STREME identifies overrepresented seed words, subsequently extended to form motif candidates. This seed-based approach reduces the search space, thereby mitigating computational complexity. A key advantage of STREME is its ability to discover motifs of different lengths in a single ST traversal. To statistically evaluate seed words enrichment in foreground sequences, STREME uses either Fisher’s exact test or the binomial distribution, depending on dataset composition (see **Supplementary File Section 1**). Once statistically significant words are identified, STREME counts approximate matches to derive PWMs. By using significant motif candidates, STREME constructs alignment profiles and iteratively refines them, assessing their statistical significance throughout the process. Profiles refinement ensures the selection of the most significant alignments, improving PWMs’ accuracy.

### 2.2 Alignment-based Motif Discovery for PWM Training: An Example with MEME

MEME [55—57] stands as a fundamental tool for motif discovery, employing an alignment-based approach to discover TFBS motifs (**Supplementary File Section 2**). Alignment-based methods construct alignment profiles by selecting motif candidate sequences from the input dataset and then evaluate them using nucleotide conservation, information content, or profile’s statistical significance [23,32]. However, finding the best alignment according to the chosen metric is computationally challenging, since it implicitly requires combining all possible subsequences from the input dataset in the alignment profile. Heuristic techniques, such as expectation maximization (EM) [55], help overcome this limitation by constructing the best alignment without exploring the entire solution space. MEME employs EM to construct PWMs while exploring the solution search space. Unlike methods using greedy approaches [66], which may focus on partial solutions, MEME ensures a more comprehensive search. For a motif of length *M,* MEME initializes an alignment profile by selecting one subsequence of length *M* from each input sequence. The initial profile is then iteratively refined using an EM. In the E-step, MEME scores each subsequence of length *M* based on their similarity to the current profile. During the M-step, MEME assigns a weight to each subsequence in the profile proportional to the scores computed in the E-step. Low-scoring subsequences are then replaced with better-fitting ones, updating the profile. The refinement process continues until convergence (i.e., the profile is no longer updated) or after a predefined number of iterations. MEME integrates various background models to assess profile significance of each constructed profile, measuring the likelihood of obtaining a profile with similar information content by chance.

## 3 Support Vector Machines-based Motif Models

SVMs have successfully addressed various biological challenges, including learning TFBSs structure [23,38]. By analyzing k-mers (DNA fragments of length *k*) frequencies in positive and negative datasets, SVM-based models train kernels to capture the structural features of binding sites. Input sequences are partitioned into *k*-mers, and their frequencies are used to create feature vectors that serve as inputs for training SVM kernels. The choice of the kernel function is critical, as it determines how SVMs represent TFs binding preferences [38]. Two primary methods are typically employed: (i) binding sites are represented as a list of *k*-mers, each with an associated weight indicating their contribution to the motif, and (ii) as a list of support vectors characterizing positive and negative datasets. One key advantage of SVM-based models over PWMs is their ability to indirectly capture higher-order dependencies between neighboring nucleotides (*k*-th order dependencies) [23,38]. Consequently, SVM-based models learn not only the binding sequence but also the context surrounding the TFBS, enabling the model to recognize complex motifs and structural patterns. For visualization purposes SVM-based models are often reduced to PWMs, obtained aligning the most informative *k*-mers. However, this conversion results in the loss of several information learned by the. Various kernels have been proposed to better capture motif structures. Early kernels, such as the spectrum kernel [67], count exact matches of all contiguous *k*-mers in the sequence data to construct feature vectors. The weighted version of the spectrum kernel [68] incorporates positional information, where the contribution of each *k*-mer to the model is weighted according to its position within the sequence. However, this approach is limited to *k*-mers up to 10 bp in length and is inefficient on large sequence datasets. More flexible kernels, such as mismatch and wildcard kernels [69,70], allow a fixed number of mismatches at specific positions within each *k*-mer, improving scalability. The dinucleotide mismatch kernel [71] further extends this idea by incorporating a first-order Markov model to capture dependencies between neighboring nucleotides. However, this method is optimized for short *k*-mers (∼10 bp), limiting its ability to characterize longer motifs. However, since binding sites generally exceed 10 bp, characterizing longer motifs using short *k*-mers may pose challenges. Additionally, using larger *k* values results in sparse feature vectors, causing the model to overfit training data. Gapped *k*-mers [72] mitigate these issues allowing gaps in non-informative or degenerate positions within the motif. This method enables the analysis of longer motifs without sacrificing scalability. Gkm-SVM [73] proposed a method to generate feature vectors for each input sequence by counting the occurrences of gapped *k*-mers in both positive and negative datasets. Feature vectors derived from gapped *k*-mers are used to train an SVM kernel, which computes the support vectors characterizing positive and negative sequence datasets. However, this process has been shown to be not scalable on datasets exceeding ∼10,000 sequences. LS-GKM [58] addresses this limitation by introducing a more efficient framework extending Gkm-SVM’s capabilities to efficiently handle large datasets. LS-GKM also provides multiple kernel functions, enhancing its flexibility in training SVM-based motif models, and offering the opportunity to learn complex sequence features contributing to TF-DNA interactions.

### 3.1 LS-GKM

LS-GKM [58] efficiently analyzes large-scale sequence datasets leveraging gapped *k*-mers to train SVM-based motif models (**Supplementary File Section 3**). By comparing datasets of bound (positive) and unbound (negative) sequences, LS-GKM generates feature vectors by counting gapped *k*-mer occurrences in both datasets. To compute gapped *k*-mer frequencies, LS-GKM builds a *k*-mer’s tree [73], whose height is *k* with a total of 4^*l*^ leaves, each corresponding to a *k*-mer. To derive feature vectors for each sequence, LS-GKM traverses the tree using a depth-first search. LS-GKM implements parallel searches on the *k*-mer tree by dividing it into smaller subtrees, processing independently each subtree. The resulting feature vectors are used to train the gapped *k*-mer kernel, which computes support vectors to distinguish between bound and unbound sequences. To ensure scalability with large datasets, LS-GKM integrates the gapped *k*-mer kernel with decomposition techniques for SVM training [58]. To compute the SVM hyperplane, the feature vectors for each sequence are mapped to the normalized frequencies of distinct gapped *k*-mers. LS-GKM offers different kernel functions to compute both linear and non-linear hyperplanes and weight the training features (**Supplementary Data Section 3**). Among these, the *gkmrbf* kernel applies a radial basis function (RBF) to the space of gapped *k*-mer frequency vectors. The center weighted gkm (*wgkm*) kernel assigns different weights to gapped *k*-mers based on their distance from the sequence center, reflecting the observation that ChIP-seq signals are often concentrated in the central regions of peaks. The *wgkmrbf* kernel combines the features of both the gkmrbf and wgkm kernels, blending their strengths for flexible and accurate modeling.

## 4 Analysis Workflow for Training and Evaluating PWM and SVM-based Motif Models

In this study, we benchmarked models’ performance in predicting TFBS by curating a diverse set of human TF ChIP-seq peak data from ENCODE. This evaluation framework incorporated synthetic and biologically feasible background data, enabling a comprehensive assessment of model robustness across different genomic contexts. Building on this foundation, we employed a systematic approach integrating model training and evaluation to assess the robustness and efficacy of PWM and SVM-based motif models in TFBS prediction. We analyzed each model’s adaptability across diverse dataset conditions, accounting for the complexities and potential noise in biological data. Our evaluation focused on key aspects of model performance, including robustness, generalizability, and reliability, under diverse training and testing scenarios, providing a comprehensive assessment of each model’s predictive capability.

### 4.1 ChIP-seq Sequence Benchmark Datasets

To benchmark models’ performance, we downloaded optimal IDR thresholded ChIP-seq peaks from ENCODE (snapshot November 2023) for 59 distinct transcription factors (TFs). To strike a balance between comprehensive TF representation and computational efficiency, we selected either a representative pair or the single available TF corresponding to each DNA-binding domain proposed in [1] (**Table 1**).

**Table 1.**
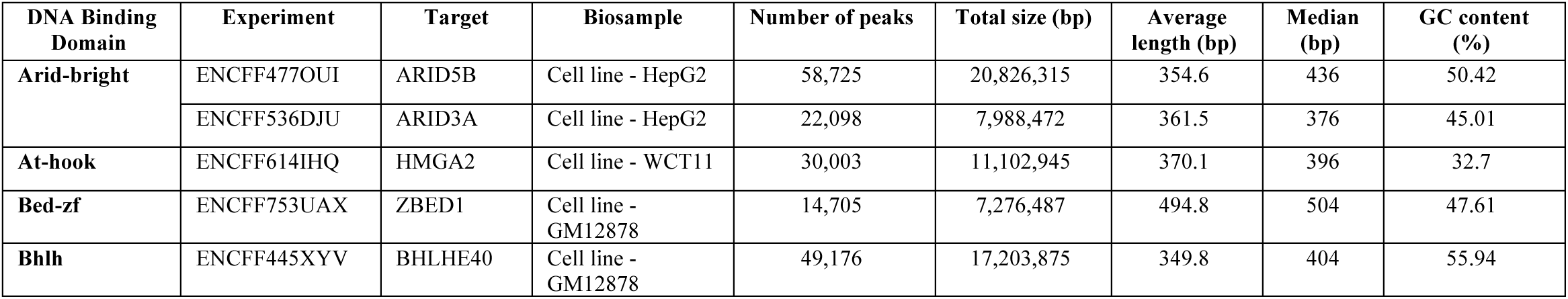
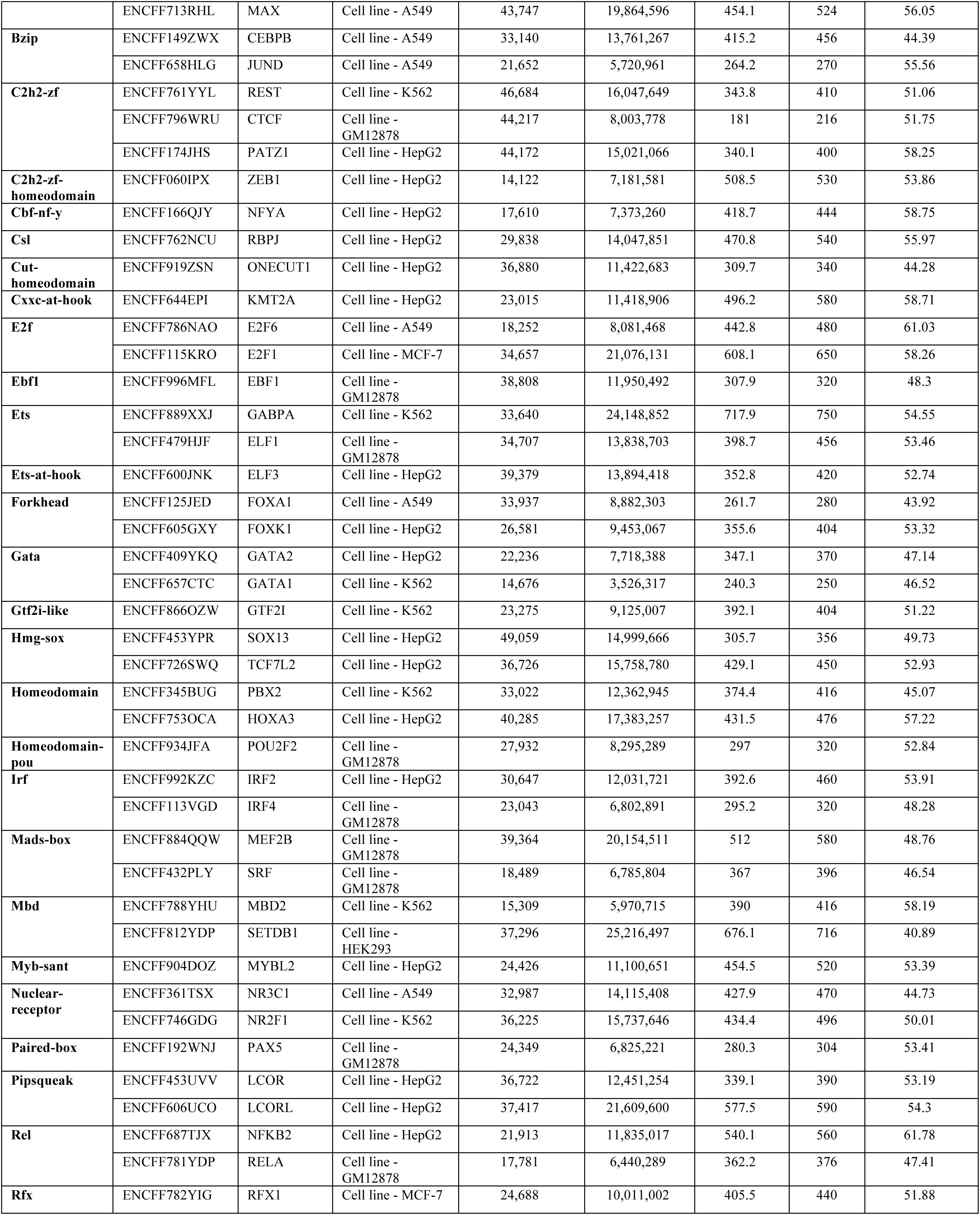
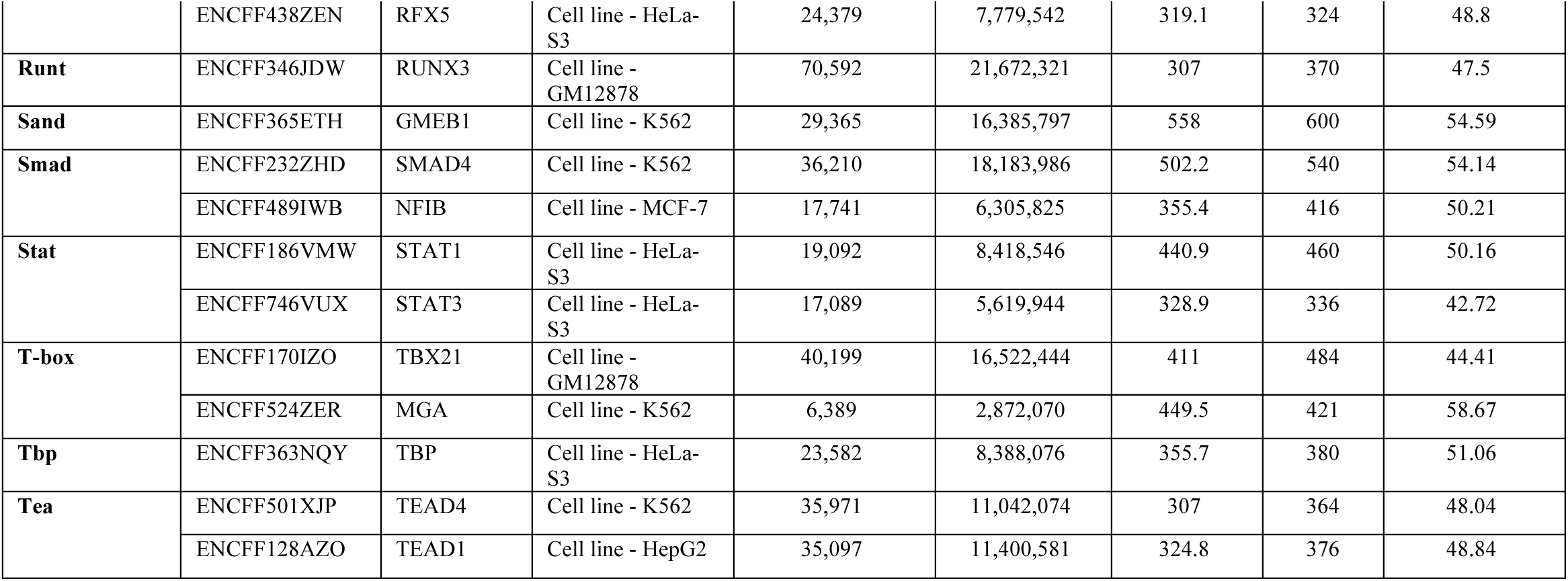
Summary statistics of the 59 ChIP-seq experiments used in the benchmark study. For each experiment, the table reports the target transcription factor, the DNA binding domain of the TF, the cell line used to perform the experiment, the number of peak sequences contained in each experiment, the total length (in bp) of peak sequences, the peaks average and median lengths, and the GC content.

For model training and performance evaluation, it is crucial to include negative sequences alongside positive ChIP-seq peak sequences. To address this, we employed two complementary strategies employing synthetic and biological background datasets. **Supplementary File Section 4** describes the procedure employed to create the background datasets. Synthetic backgrounds were generated by shuffling the nucleotides of positive sequences, while preserving first-order dependencies (**Figure 1A**). This approach ensures that nucleotide pairs remain intact, thereby creating more biologically plausible backgrounds [74]. For biologically feasible background data, we used a comprehensive collection of DNase I hypersensitive sites (DNase-seq) derived from various tissue types [75]. To ensure that background and ChIP-seq sequences were distinct, we selected DNase-seq regions not overlapping, even partially, any of the peaks in the corresponding ChIP-seq dataset (**Figure 1A**). To ensure consistent genomic contexts, we restricted DNase-seq regions to match chromosomes on positive peaks. To split positive and negative datasets into training and testing sets, we allocated all sequences on chromosome 2 to the test sets, leaving the rest for model training. This chromosome-based split ensured that the test dataset was independent from training data.

**Figure 1.**
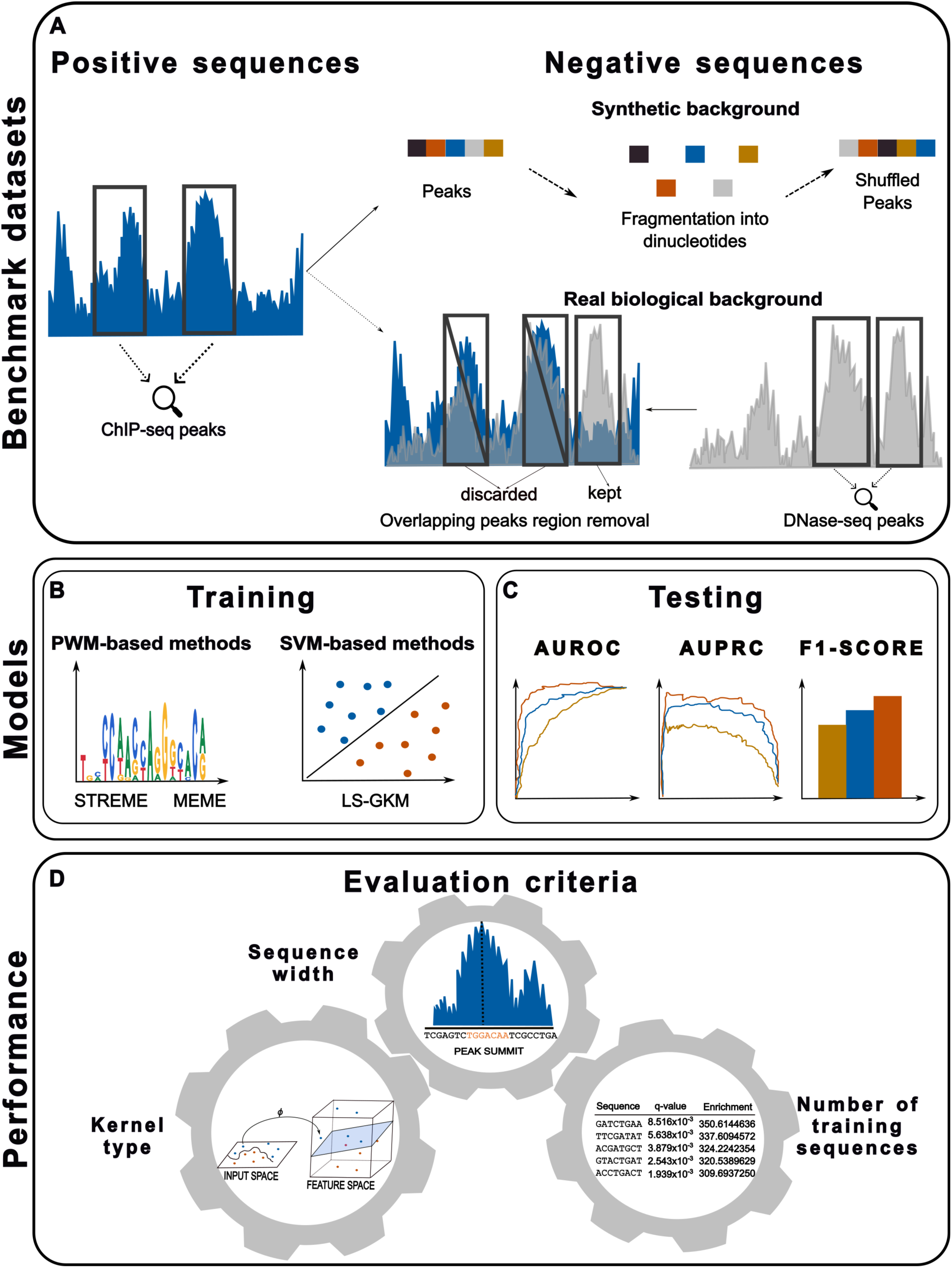
Analysis workflow to train and evaluate PWM and SVM-based motif models. **(A)** Positive sequences are derived from ChIP-seq peaks, representing genomic regions enriched for transcription factor binding. Negative sequences are generated using two approaches: (i) synthetic construction by shuffling dinucleotides from the positive sequences to preserve local sequence composition, and (ii) selection from DNase-seq hypersensitive sites that do not overlap with ChIP-seq peaks, providing real-world background data. These datasets reflect diverse biological and computational scenarios for model evaluation. **(B)** Motif models are trained using PWM-based approaches, including STREME and MEME, as well as SVM-based methods like LS-GKM. PWMs capture nucleotide frequencies at each motif position, while SVM-based models leverage kernel functions to identify complex sequence patterns and positional dependencies. **(C)** The performance of each model is assessed using multiple metrics, including Area Under the Receiver Operating Characteristic (AUROC), Area Under the Precision-Recall Curve (AUPRC), and F1-Score. These metrics measure a model’s ability to accurately distinguish true binding motifs from background sequences across varying conditions. **(D)** Key factors influencing model performance are systematically explored, including dataset size, sequence width, and kernel type for SVM-based models. This analysis identifies optimal configurations for each model and highlights their strengths and limitations across different data properties and biological contexts.

Synthetic and biological background datasets introduced two testing scenarios, enabling a nuanced evaluation of models’ performance. Synthetic background data test the models’ ability to distinguish true TFBS from artificially generated negative sequences. This approach creates a controlled environment emphasizing motif-recognize capability, while excluding the influence of complex genomic features. Conversely, biologically feasible background data simulate genomic regions with accessible DNA but not necessarily bound by the investigated TF. In this scenario, the challenge for the model is to distinguish between TF binding events from regions of open chromatin that lack TF binding. To further assess models’ predictive ability in identifying TFBS within large genomic contexts, we constructed an additional test dataset with a 10x higher proportion of DNase-seq sequences compared to positive sequences. This scenario provides insights into models’ robustness under realistic genomic conditions, where biologically active but non-binding regions are prevalent, resulting in class imbalance.

### 4.2 Models Training and Testing

Training was conducted independently for SVM and PWM models, with MEME/STREME used to train PWMs and LS-GKM for generating SVM-based motifs (**Figure 1B**). **Supplementary File Section 5** details the training procedure. Each model was evaluated across four test conditions designed to assess models’ robustness, adaptability, and susceptibility to overfitting. (i) Training and testing on synthetic negative datasets provide a controlled environment to establish a baseline for model performance in the presence of simple, random noise. (ii) Training and testing on DNase-seq background data replicate a realistic biological setting, testing each model’s ability to function under complex, biologically relevant background conditions where TF binding might plausibly occur [76,77]. (iii) Training on synthetic negatives and testing on DNase-seq data is a technique often employed [77—79] and introduce a background complexity shift evaluating whether models trained on simpler data could adapt effectively to biologically complex contexts. (iv) Training on DNase-seq background sequences and testing on synthetic data reverse the previous scenario, examining whether models trained in a complex biological context could retain accuracy and generalizability when evaluated against a simpler background. For tests involving biological background data two scenarios were evaluated: (i) a balanced setup with a 1:1 ratio of positive to negative sequences, and (ii) an imbalanced setup with a 1:10 ratio. The latter simulated real-world conditions, where unbound genomic regions outnumber bound regions. These test conditions provide a comprehensive assessment of each model’s generalizability, resilience to varying background complexities, and suitability for TFBS prediction in realistic genomic settings.

After training, models were employed to score each test sequence and assess the likelihood of TF binding. For SVM-based models, scores were generated using LS-GKM’s *gkmpredict* function [58], while for PWMs generated by MEME/STREME, scoring was performed using FIMO [80] (see **Supplementary File Section 6** for details). Following sequence scoring with the trained models, we evaluated their predictive performance across different testing scenarios using multiple metrics (**Figure 1C**).

### 4.3 Models evaluation criteria: the Impact of Dataset Size, Sequence Width, and Kernel Functions

To comprehensively assess the predictive performance and robustness of SVM and PWM models, we performed models training across different experimental conditions (**Figure 1D**). Each setting was chosen to explore key factors that may influence model accuracy and generalizability, such as the number of training sequences, sequences width, and the choice of kernel function, for SVM models. The first analyzed factor was training dataset size (**Figure 1D**). Models were trained on datasets of varying sizes (500, 1000, 2000, 5000, 10000, and the full dataset), prioritizing the most reliable sequences (peaks sorted by *q*-value and ChIP enrichment score). This analysis aimed to determine the effect of dataset size on model performance. While smaller datasets can reduce computational requirements, they may not capture the full binding sites complexity, potentially impacting models’ perfromance [46,47]. On the other hand, larger datasets contain more information, but they may introduce noise from less informative sequences [81]. Next, the sequence width was systematically varied to evaluate its influence on model performance (**Figure 1D**). Sequences of different lengths (50, 100, 150, 200, and full-length) were extracted around the ChIP-seq peak summit and used for training and evaluation. This analysis illustrates how training sequences’ length influences the surrounding genomic context that may either support or interfere with accurate motif models’ training [49]. Shorter sequences may focus the model on the core binding site, reducing noise from surrounding regions, but could also omit important flanking information that might contribute to TF’s binding specificity [82]. On the other hand, longer sequences include flanking regions potentially introducing noise but provide a richer context for motif training. Lastly, for SVM-based models, we compared different kernel functions available in LS-GKM (**Supplementary File Section 3**): these included the gapped *k*-mer kernel, *k*-mer frequency counts, the standard *gkm* kernel, the *gkm* kernel combined with a radial basis function (RBF), and the center-weighted variants of both the *gkm* and *gkm*-RBF kernels (**Figure 1D**). Additionally, configurations better performing by tool and by individual experiment were evaluated to provide a broader perspective on model performance.

### 4.4 Models Performance Metrics

In our study, we defined true positives (TP) as test sequences from ChIP-seq datasets that were correctly predicted by the model to be bound by the investigated TF. False positives (FP) were sequences from the negative test dataset that the model incorrectly predicted as bound by the TF. Conversely, sequences in the positive dataset that were predicted as unbound were labeled as false negatives (FN). Given the difficulty in accurately determining the total number of negative binding sites for each TF across the genome [83], we opted to exclude performance metrics that depend on true negative (TN) counts [84]. As a result, we evaluated model performance using precision 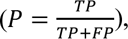 recall 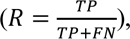 and F1-score 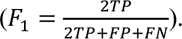 Additionally, to further assess model performance, we computed the area under the precision-recall curve (AUPRC) and the area under the receiver operating characteristic curve (AUROC) for each model and test dataset (**Figure 1C**).

## 5 Benchmarking PWM and SVM Motif Models Performance

In this section, we compare the performance of models trained using MEME, STREME, and LS-GKM by analyzing key performance metrics, such as AUPRC, AUROC, and F1-score under multiple training and testing scenarios. We explore how these metrics evolve as experimental settings change, leveraging both synthetic and biologically feasible background training data.

By analyzing how varying training dataset size and background data affects model performance, we observed distinct trends across methods. Since MEME ignores background data to train PWMs, its performance remains stable regardless of background changes (**Figure 2A** and **B**). Across 59 datasets, MEME performs best at distinguishing positive (ChIP-seq) sequences from synthetic negatives when trained on the most reliable 5,000 sequences (AUPRC = 0.7). Interestingly, when distinguishing positives from biological negatives (1:1 ratio), MEME performs better when trained on the top 1,000 sequences (AUPRC = 0.72). However, performance drops when tested on the needle-in-the-haystack scenario (1:10 positive-negative ratio, AUPRC = 0.28), but 1,000 sequences still yield the best results. Similar trends are observed for AUROC (**Supplementary Figure 1A and B**), while MEME performs best at 5,000 sequences on all three test scenarios in terms of F1-score (**Supplementary Figure 2A and B**). STREME shows different trends depending on the training background (**Figure 2C** and **D**). Using synthetic background data, when tested on distinguishing positive from synthetic negatives, STREME achieves the best results at 10,000 sequences (AUPRC = 0.72). Even when tested on biological negatives, STREME remains competitive (AUPRC = 0.77) performing best at 10,000 sequences. However, in the needle-in-the-haystack scenario, performance declines sharply (AUPRC = 0.35). AUROC and F1-score follow similar trends (**Supplementary Figures 1C and 2C**). When trained using biological background, STREME’s performance on synthetic test data generally decreases, but it improves significantly on biologically feasible negatives (AUPRC = 0.82 for 1:1 and AUPRC = 0.43 for 1:10), with dataset sizes of 10,000 and full yielding the best performance. AUROC and F1-score confirm that larger training sets enhance STREME’s performance in this scenario (**Supplementary Figures 1D and 2D**). SVM-based models’ performance is more influenced by the choice of background data (**Figure 2E** and **F**). When trained using synthetic backgrounds, they perform best on synthetic test data (AUPRC = 0.97 at size full), whereas training using biological negatives improves performance on biologically feasible test sets (highest AUPRC = 0.96 for 1:1 and AUPRC = 0.8 for 1:10 at size full). These models benefit from larger training datasets, reaching peak performance when trained on full datasets. SVMs exhibit robustness even in the needle-in-the-haystack scenario, particularly when trained using biological negatives (AUPRC = 0.8 vs. AUPRC = 0.3 for synthetic-trained models). AUROC and F1-score analyses confirm that larger training sets and careful choice of background training data enhance SVMs performance (**Supplementary Figures 1E-F** and **2E-F**).

**Figure 2.**
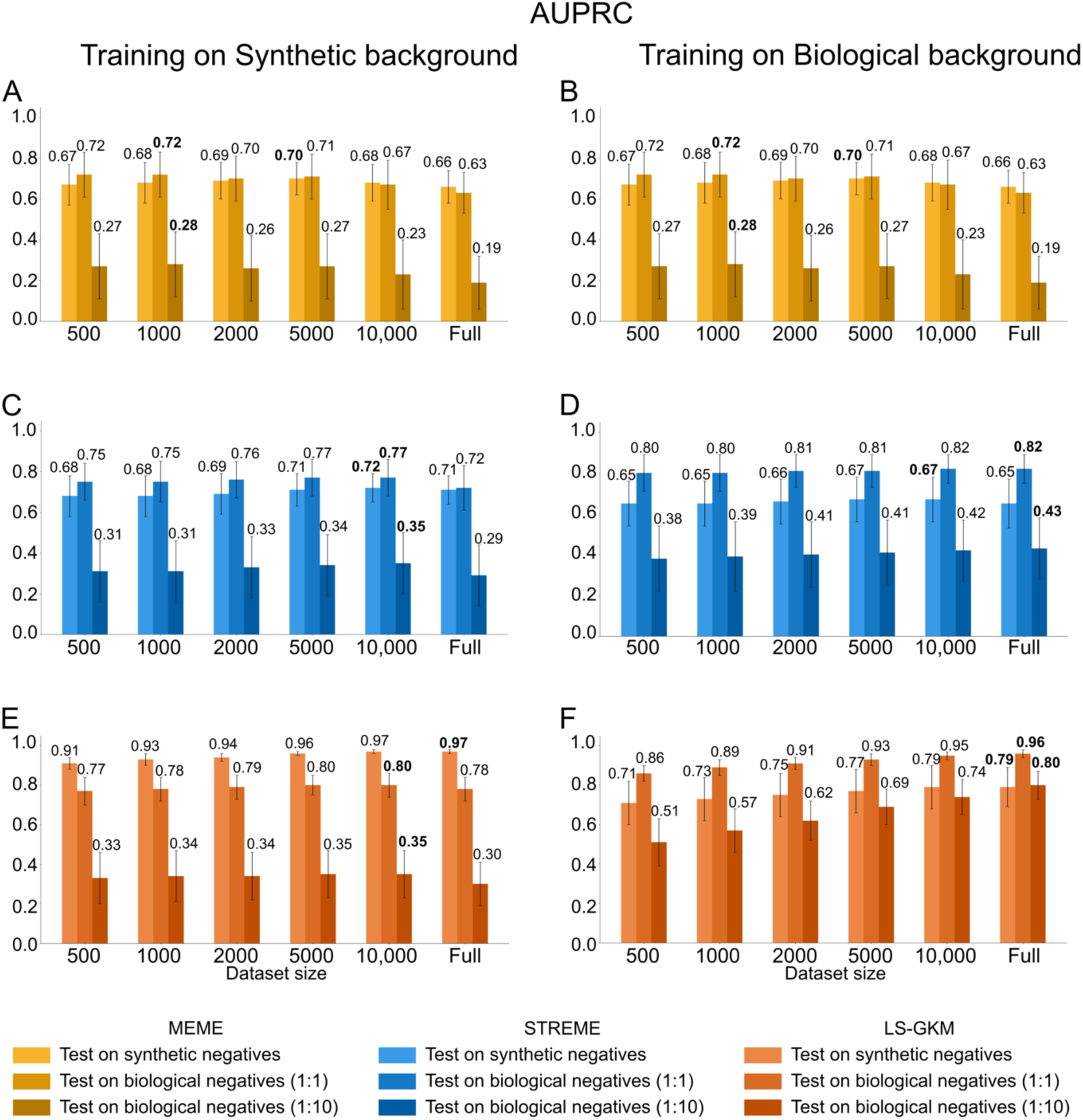
Average AUPRC values from models trained across 59 datasets with varying background and dataset sizes. **(A,B)** AUPRC performance of MEME’s PWMs trained on synthetic **(A)** or biologically feasible **(B)** background data, tested on synthetic negative sequences and biological negatives (1:1 and 1:10 positive-negative ratios). **(C, D)** AUPRC performance of STREME’s PWMs trained on synthetic **(C)** or biologically feasible **(D)** background data, evaluated under the same testing conditions. **(E, F)** AUPRC performance of SVM-based models trained on synthetic **(E)** or biological **(F)** background data, tested similarly. Labels in bold mark the best performing training dataset size on each testing scenario.

Examining how training sequence width affects model performance revealed distinct trends across methods. MEME generally performs best when trained on narrow sequences centered around peak summits (**Figure 3A** and **B**), peaking at 100 bp when tested on synthetic (AUPRC = 0.71) and biological negatives (1:1 ratio; AUPRC

**Figure 3.**
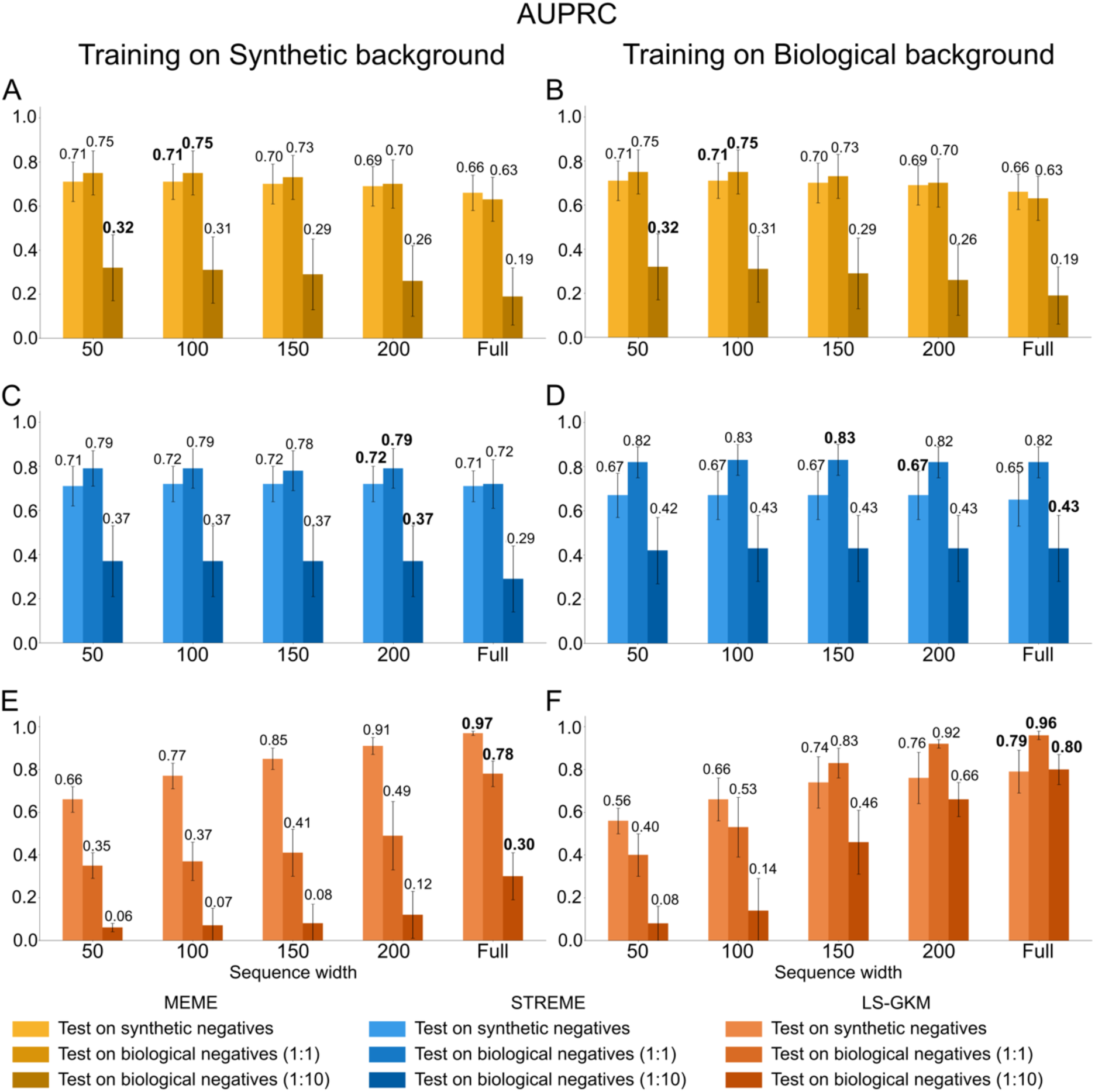
Average AUPRC values from models trained across 59 datasets with varying background and sequence widths. **(A,B)** AUPRC performance of MEME’s PWMs trained on synthetic **(A)** or biologically feasible **(B)** background data, tested on synthetic negative sequences and biological negatives (1:1 and 1:10 positive-negative ratios). **(C, D)** AUPRC performance of STREME’s PWMs trained on synthetic **(C)** or biologically feasible **(D)** background data, evaluated under the same testing conditions. **(E, F)** AUPRC performance of SVM-based models trained on synthetic **(E)** or biological **(F)** background data, tested similarly. Labels in bold mark the best performing training sequence width on each testing scenario.

= 0.75). For the needle-in-the-haystack scenario, MEME achieves the best performance at 50 bp sequences. AUROC and F1-score analyses confirm that MEME benefits from shorter sequences (**Supplementary Figures 3A-B** and **4A-B**). STREME, instead, displays different performance depending on the background used during training (**Figure 3C** and **D**). STREME performs best at 200 bp when tested on synthetic negatives, whether trained on synthetic (AUPRC = 0.72) or biological backgrounds (AUPRC = 0.67). On biological negatives (1:1 ratio), optimal performance occurs at 200 bp (AUPRC = 0.79) when trained on synthetic background and at 150 bp (AUPRC = 0.83) when trained on biological feasible background. In the needle-in-the-haystack scenario, STREME benefits from using biological background during training, with full-length sequences yielding the best performance (AUPRC = 0.43 vs. AUPRC = 0.37 for synthetic-trained models). AUROC and F1-score trends align with these findings (**Supplementary Figures 3C-D** and **4C-D**), indicating that while STREME benefits from larger training sets than MEME, the method is still sensitive to noise introduced by sequences flanking TFBSs. SVMs display susceptibility to sequence width (**Figure 3E** and **F**). In fact, they achieve performance comparable to those achieved by MEME and STREME only when trained on full-length sequences when trained using synthetic background and 150 bp using biological background. The best results occur when trained using biological background sequences with full-length sequences and tested on biologically feasible negatives (AUPRC = 0.96 and 0.8, respectively). AUROC and F1-score analyses confirm these observations, further highlighting the advantages of using longer training sequences for SVM-based models (**Supplementary Figures 3E-F** and **4E-F**).

We also evaluated the impact of kernel functions on SVM-based model performance (**Figure 4A** and **B**). When trained and tested using synthetic background data (**Figure 4A**), gapped *k*-mer, *gkm*, and *gkmrbf* kernels outperform the classical *k*-mer counting and weighted kernels across all test scenarios. When tested on biological negatives, instead, the weighted *gkm* and *gkmrbf* kernels achieve the best performance (AUPRC = 0.78 for 1:1 ratio, 0.3 for 1:10 ratio). Similarly, when trained using biologically feasible background sequences (**Figure 4B**), weighted kernels consistently outperform the other kernels on all test scenarios, indicating that positional weighting enhances model robustness particularly when analyzing biologically feasible data (AUPRC = 0.8 for both *wgkm* and *wgkmrbf*). AUROC and F1-score analyses confirm these trends (**Supplementary Figures 5A-B** and **6A-B**).

**Figure 4.**
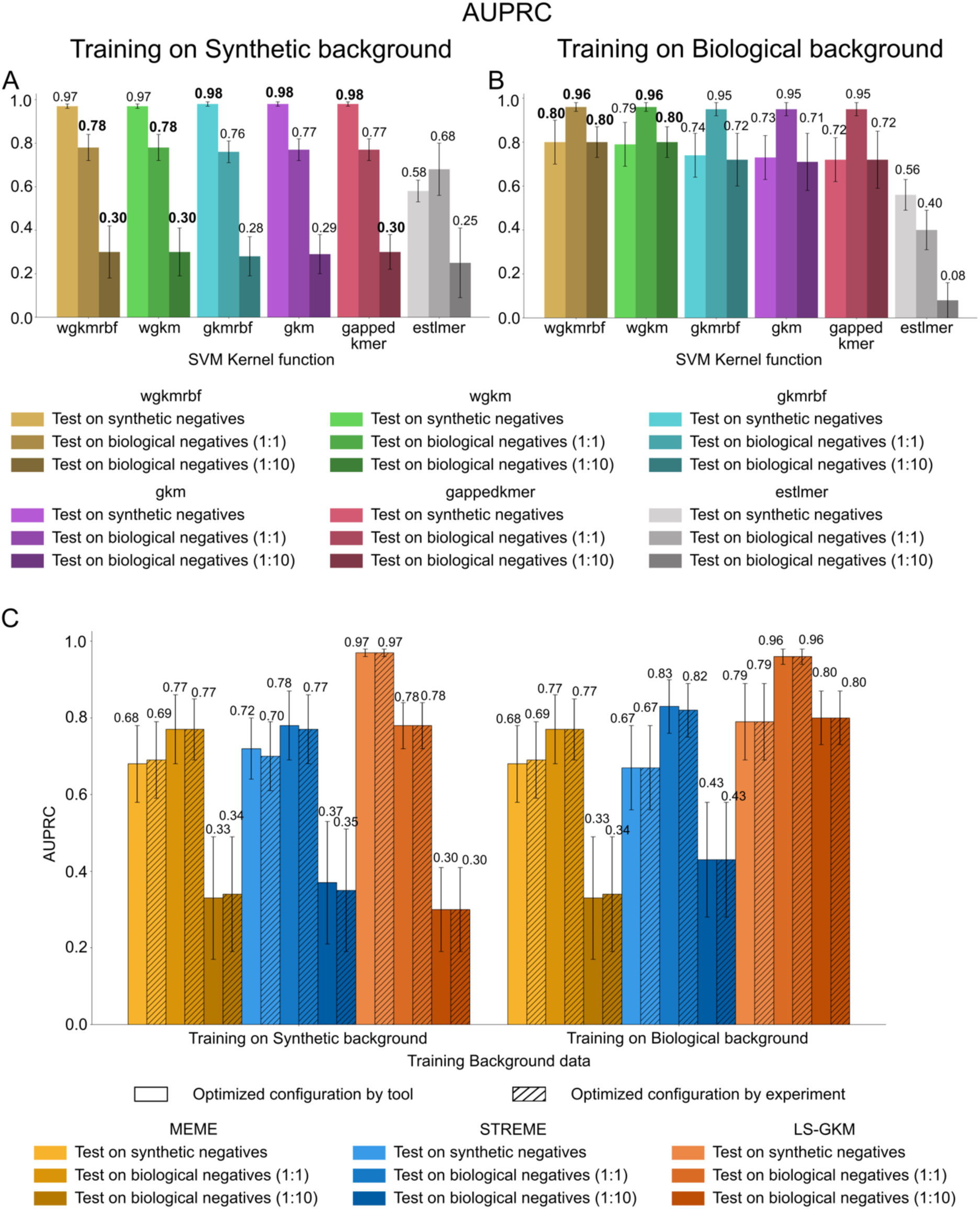
Average AUPRC values for SVM-based models trained across 59 datasets, comparing different LS-GKM kernel functions and optimized training configurations. **(A,B)** Performance of kernel functions available in LS-GKM when trained on synthetic **(A)** and biologically feasible **(B)** background sequences. Labels in bold mark the best performing training kernel function on each testing scenario. **(C)** Comparison of model performance using two optimization strategies: a globally optimized tool-specific configuration versus per-experiment fine-tuning.

To assess the impact of configuration optimization on model performance, we applied two optimization strategies: (i) selecting the best-performing dataset size, sequence width, and kernel (for SVMs) per tool across all experiments, and (ii) selecting the best configuration individually for each dataset and tool (**Figure 4C**). By analyzing average performance across AUPRC, AUROC, and F1-score (**Figure 4C** and **Supplementary Figures 5C** and **6C**) we generally observed an improvement of performance compared to those obtained in non-optimized scenarios. However, we observed no significant differences between the two optimization approaches across training scenarios (**Supplementary Table 1**). This suggests that a globally optimized, tool-specific configuration is sufficient for robust performance, eliminating the need for fine-tuning on a per-experiment basis.

To investigate the sequence patterns driving models’ prediction, we investigated the positional distribution of motif scores on positive test sequences for the best- and worst-performing TFs for each method (**Figure 5** and **Supplementary Figure 7**). For each TF, models were applied to score positive test sequences (from test sets) at each position. Scores measure the likelihood that the binding motif of the investigated TF occurs at a specific position within the sequence.

**Figure 5.**
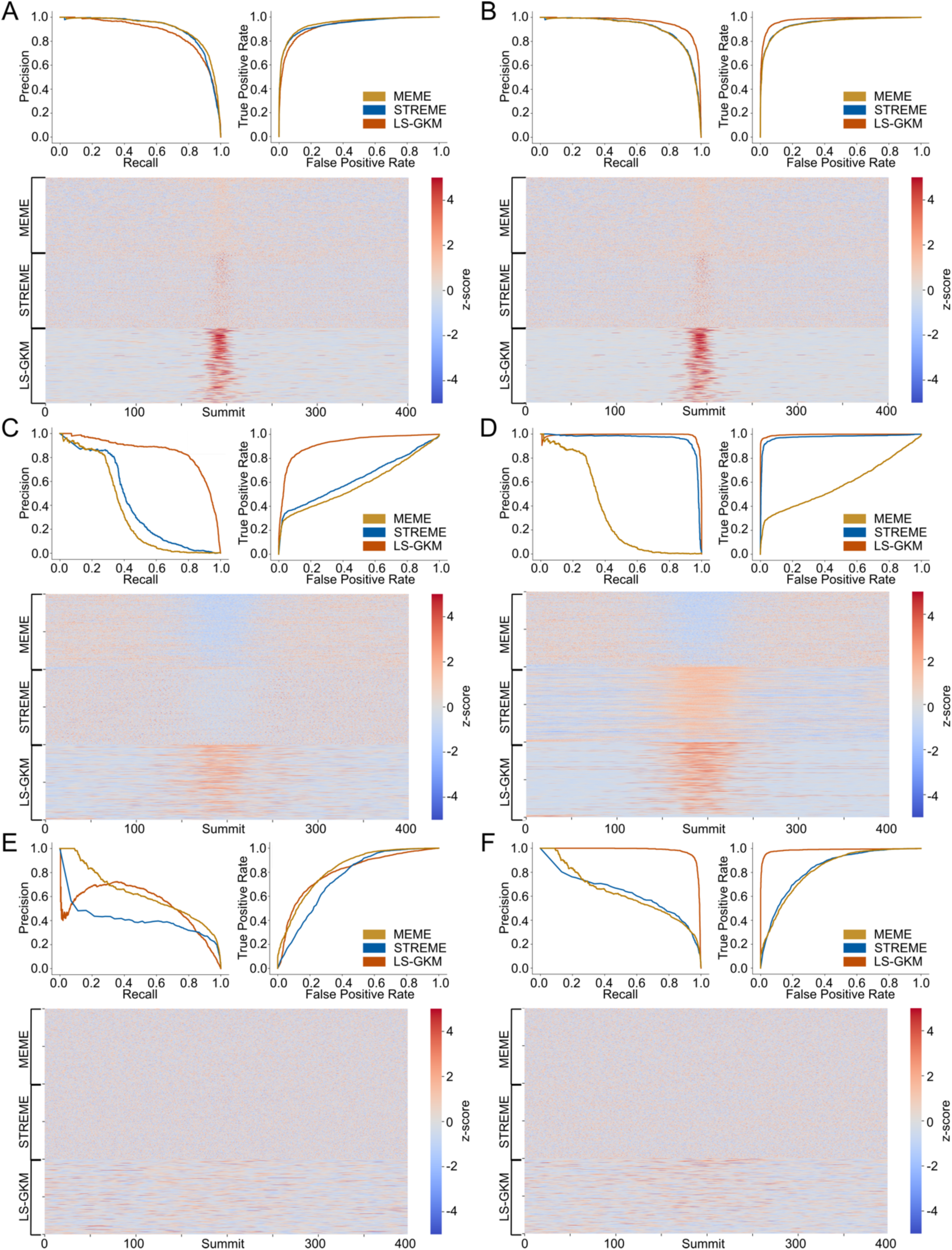
Comparison of transcription factor binding predictions by MEME, STREME, and LS-GKM trained on synthetic (left) and biological (right) background sequences. The panels present results for the top-performing TFs of each tool: **(A, B**) CTCF for MEME (experiment ENCFF796WRU), (**C, D**) HMGA2 for STREME (experiment ENCFF614IHQ), and (**E, F**) SETDB1 for LS-GKM (experiment ENCFF812YDP). Heatmaps illustrate the positional distribution of motif scores across test sequences, highlighting motif enrichment patterns near peak summits. ROC (top left) and precision-recall (top right) curves summarize predictive performance.

MEME performed best in predicting CTCF [85] (experiment ENCFF796WRU) binding (**Figure 5A-B**), showing motif occurrences near peak summits even when trained using synthetic background sequences.STREME exhibited a similar but stronger trend. LS-GKM displayed a sharper signal localization, aligning with its higher AUPRC compared to PWMs (**Figure 5A**). Training models using biologically feasible background sequences (**Figure 5B**) yielded a minor improvement for LS-GKM, suggesting that it benefits from training with biologically relevant backgrounds but remains robust across training conditions. For MEME and STREME PWMs the improvement was more marked. Precision-recall (PRC) and receiver operating characteristic (ROC) curves further support these findings, confirming SVM’s predictive advantage.

STREME excelled in predicting HMGA2 [86] (experiment ENCFF614IHQ) binding (**Figure 5C-D**), particularly when trained using biological negatives. In this scenario, STREME’s PWM showed a strong motif signal around the peak summit (**Figure 5D**). However, the signal was less pronounced than with LS-GKM, which maintained the highest signal concentration around the peak summit. MEME, instead, displayed a dispersed signal instead. When trained employing synthetic background sequences, MEME and STREME performed near-randomly (**Figure 5C**), failing to detect binding signals. Similarly, LS-GKM’s SVM model performed less effectively in this scenario. PRCs and ROCs reinforce these trends, highlighting LS-GKM’s consistent advantage and STREME’s dependence on training background sequences.

LS-GKM achieved its highest performance predicting SETDB1 [87] (experiment ENCFF812YDP) binding (**Figure 5E-F**), outperforming MEME and STREME when trained using biologically feasible background sequences (**Figure 5F**). However, its predictions exhibited broader signal distribution compared to the previous TFs, potentially indicating that the model captured secondary, functionally relevant sequences beyond the binding site itself. This suggests that LS-GKM may incorporate additional regulatory elements into its predictions, a hypothesis supported by PRC and ROC curves, which reflect the overall challenges in predicting SETDB1 binding.

An analysis of the experiments with the weakest predictive performance revealed consistent trends across the three tools. MEME showed the lowest AUPRC when predicting MYBL2 [88] (experiment ENCFF904DOZ) binding (**Supplementary Figure 7A-B**). Heatmaps show that none of the models, including the SVM-based model, captured clear positional enrichment when trained using synthetic background data. Similar results were observed training models with biological negatives, where MEME and STREME failed to display motif aggregation (**Supplementary Figure 7A**). However, LS-GKM exhibited slight improvement, with stronger signal concentration near the peak summit (**Supplementary Figure 7B**) and increased AUPRC and AUROC.

STREME performed worst when predicting SOX13 [89] (experiment ENCFF453YPR) binding (**Supplementary Figure 7C-D**). When trained using synthetic background sequences, it showed no clear positional weighting (**Supplementary Figure 7C**), while training with biological background data led to a light improvement (**Supplementary Figure 7D**. LS-GKM, in contrast, maintained strong performance across both training conditions, consistently assigning higher weights to regions near the peak summit. MEME behaved randomly in both conditions.

LS-GKM struggled most with TEAD4 [90] (experiment ENCFF501XJP) binding (**Supplementary Figure 7E-F**). Heatmaps reveal that PWMs failed to learn informative patterns, showing no motif enrichment near the peak summit. However, LS-GKM still outperformed MEME and STREME, displaying robustness even in challenging prediction scenarios. This suggests LS-GKM may better handle complex binding grammars, albeit with some limitations.

Analyzing the tool’s scaling behaviors, they displayed different trends as dataset size and sequence width increase (**Supplementary Tables 2 and 3**). LS-GKM experiences a sharp rise in processing time with larger datasets and wider sequences, leading to high computational costs. MEME also slows considerably under these conditions. In contrast, STREME is the fastest for smaller datasets but becomes increasingly sensitive to input size, eventually surpassing MEME in processing time. Among LS-GKM’s kernel functions, the gkm kernel is the most efficient, while the classical k-mer counting kernel (estlmer) is the slowest. Computational performances were measured on a Linux PC with an AMD Ryzen Threadripper 3970X CPU with 32 cores and 64 GB of RAM.

## 6 Creating a comprehensive SVM Models’ Collection

SVM-based motif models generally outperform PWMs in predicting TF binding, particularly when evaluated on biologically feasible datasets containing positive and negative test sequences. SVM-based models exhibit robustness against dataset imbalance, delivering strong performance even when negative sequences significantly outnumber positive ones.

Despite their predictive performance, SVM-based models have seen limited adoption. This can be attributed to several factors. One key challenge is their interpretability: unlike PWMs, which can be visualized as motif logos that intuitively convey the learned features, SVM-based models lack a straightforward method for interpreting what the model has learned. Additionally, the adoption of SVM-based models is hindered by the absence of comprehensive databases of precomputed models. In contrast, PWMs are readily available in well-established repositories such as JASPAR or HOCOMOCO [63,64].

To address this gap, we developed what is, to our knowledge, the first comprehensive database of SVM-based motif models trained on human ChIP-seq data from ENCODE (snapshot December 2024). For this aim, we selected high-quality human ChIP-seq datasets (IDR thresholded peaks) mapped on hg38 genome assembly belonging to 176 different tissues and cell lines (see **Supplementary Table 4**). Each dataset was split into separate training and test sets to ensure rigorous model evaluation.

Since in our analyses we demonstrated SVM-based models better perform when trained using DNase-seq sequences as negative data, compared to shuffled sequences, we employed DNase-seq data as the background for both training and testing. Following a similar approach to our earlier analyses, we allocated all features mapped to chromosome 2 as test data, while features from all other chromosomes were used for training. In terms of preprocessing, we observed that SVM-based models achieved optimal performance when using the complete sequence dataset, including full-length peak sequences, without additional truncation or peaks trimming. Therefore, we retained the raw sequence information for training. Furthermore, based on our evaluations, the centered weighted gapped *k*-mer kernel yielded the best predictive performance on average. Thus, we utilized this kernel to train models.

For each model, we systematically evaluated its predictive performance on the test dataset using multiple metrics (AUPRC, AUROC, and F1-score). These assessments provided insights into the models’ accuracy and reliability (see **Supplementary File Section 7**). Our pipeline ultimately produced 2,393 SVM-based motif models, each optimized for predicting TF binding (see **Supplementary Table 4**). This models’ collection represents a valuable resource for the community, offering a new avenue for investigating TF binding with high predictive accuracy and biological relevance.

## 7 Discussion

In this study, we performed a comprehensive benchmark to compare the performance of PWMs and SVM-based motif models in encoding TFBSs. To thoroughly evaluate the models at both theoretical and practical levels, we designed multiple test scenarios. Specifically, we used synthetic background data to measure the theoretical predictive ability of the models and DNase-seq data as a background to provide a more realistic assessment of their performance in biologically relevant contexts.

Our results demonstrate that SVM-based models generally outperform PWMs in most scenarios, particularly when tested on imbalanced datasets. However, we also observed that PWMs can still serve as the preferred model under certain conditions. **Table 2** discusses the better performing models based on input dataset characteristics.

**Table 2.** Model Selection Guide Based on Dataset Characteristics. This table provides guidance on selecting the better model—PWM (MEME or STREME), or SVM (LS-GKM)— based on key dataset attributes, including dataset size, sequence width, and background training data type. The column **Dataset Size** refers to the number of sequences in the training dataset (Small: fewer than 5,000 sequences; Large: more than 5,000 sequences). **Sequence Width** indicates the length of each sequence in the training dataset (Narrow: less than 200 bp wide; Long: more than 200 bp wide). **Background Type** describes the type of background data used for training. The column **Recommended Model** highlights the model expected to perform best for each combination of characteristics, with a rationale provided for each recommendation.

| Dataset Size | Sequence Width | Background Type | Recommended Model | Rationale |
| --- | --- | --- | --- | --- |
| Small | Narrow | Synthetic | PWM (MEME) | MEME performs better on small datasets of narrow sequences, especially when background data is unavailable, such as in SELEX or PBM experiments, where sequences lack genomic context (**Figure 2A** and **3A**). Since MEME does not require background data to train PWM models, it handles synthetic backgrounds better, making it ideal for cases where no biologically meaningful background can be recovered. In contrast, STREME benefits from real biological background data (e.g., DNase-seq) and performs well when genomic context matters, such as in ChIP-exo experiments (**Figure 2D** and **3D**). When meaningful background sequences are available, STREME is the better choice. |
|  |  | Biologically Feasible | PWM (STREME) |  |
| Large | Narrow | Synthetic | PWM (MEME) | MEME performs well on large datasets of narrow sequences, particularly when background data is unavailable and synthetic backgrounds need to be computed (**Figure 2A** and **3A**). Since MEME does not require background data to train PWM models, it is better suited for cases where only synthetic background data may be used, avoiding potential biases from biologically implausible controls. STREME, in contrast, excels with large datasets of narrow sequences, such as ChIP-exo data (**Figure 2D** and **3D**). Its ability to integrate biologically feasible background data (e.g., DNase-seq) enhances its ability to distinguish bound from unbound sequences. |
|  |  | Biologically Feasible | PWM (STREME) |  |
| Small | Long | Synthetic | SVM (LS-GKM) | LS-GKM SVM-based models performs well on small datasets with long sequences, particularly when trained on biologically feasible background data, making it suitable for high-quality but limited ChIP-seq data where biological context enhances models' performance (**Figure 2F**). SVMs leverage the amount of information carried by long sequences. When biological background is unavailable, LS-GKM remains applicable using synthetic background data (**Figure 2E**). However, STREME PWMs, while slightly outperformed by LS-GKM in this synthetic background setting, remains competitive (**Figure 2C**). |
|  |  | Biologically Feasible | SVM (LS-GKM) |  |
| Large | Long | Synthetic | SVM (LS-GKM) | LS-GKM performs well on large datasets with long sequences, particularly when trained on biological background data. This makes it well-suited for ChIP-seq, where both binding sites and regulatory flanking regions contribute to signal (**Figure 2F** and **3F**). Even when only synthetic background data is available, LS-GKM remains effective, performing strongly in this setting (**Figure 2E** and **3E**). |
|  |  | Biologically Feasible | SVM (LS-GKM) |  |

PWMs trained using alignment-based methods like MEME are particularly valuable for small datasets with highly reliable bound sequences. These methods do not require background data for training, making them ideal for scenarios with high-confidence positive data and where background datasets cannot be recovered or computed, such as PBM or SELEX [13,14] experiments (**Table 2**).

PWM training methods based on enumeration, such as STREME, are also well-suited for small, reliable datasets and can handle larger datasets more effectively than alignment-based methods (**Table 2**). Unlike MEME, enumerative methods can employ background sequences during training. Our analysis shows that employing biologically feasible background data, such as DNase, rather than synthetic negative sequences, significantly improves their ability to distinguish between bound and unbound sequences. However, we observed that the performance of STREME is influenced by the input dataset size and the length of training sequences. Since PWMs focus on learning the binding site itself and not the surrounding sequence context [91,92], longer sequences can introduce noise, reducing predictive performance. Consequently, STREME is particularly effective for large datasets with narrow sequences, such as those generated by ChIP-exo [24].

When working with large datasets containing long sequences, as is common in ChIP-seq experiments, SVM-based models appear better suited. However, these models incur a significant increase in computational time for larger datasets (**Table 2**). These models not only learn TFBSs but also capture the surrounding sequence context, making them more robust and better at tolerating noise from flanking regions. This ability to incorporate complex features contributes to their performance, particularly when tested on imbalanced datasets, where the negative class is substantially larger than the positive class.

However, SVM-based models lack interpretability, making it challenging to determine which sequence features the models learned [23,93,94]. In scenarios where understanding the learned features is crucial, PWMs remain the preferred choice due to their simplicity and straightforward visualization. Nevertheless, analyzing SVM weight assignments to *k*-mers suggest that these models possibly recognize secondary regulatory sequences beside motifs. Furthermore, SVM training often requires significantly more time compared to PWM training methods like STREME or MEME, making PWMs a more practical option for time-sensitive applications.

In this study, we selected evaluation metrics that do not rely on assumptions about true negative sequences. Accurately identifying sequences that are unbound by a TF in real genomic datasets is inherently challenging. For instance, sequences classified as negatives, such as DNase-seq peaks that do not overlap with ChIP-seq peaks, may remain unbound under the specific experimental conditions tested. However, these same sequences might exhibit TF binding activity under alternative environmental conditions, or in different cell types or tissues [95—97].

This ambiguity underscores the importance of prioritizing metrics that do not explicitly depend on the definition of true negatives when assessing model performance, such as AUPRC. By doing so, we mitigate potential biases introduced by incorrect or context-dependent classification of negatives.

Additionally, we emphasize the value of metrics that penalize false negatives, as the identification of bound sequences is often supported by high-confidence experimental data, such as ChIP-seq, that validate TF binding. The confidence in positive sequences contrasts sharply with the uncertainties associated with negatives, making it essential to minimize errors in identifying true binding events. False negatives can have significant downstream consequences, particularly in applications such as regulatory network reconstruction [98,99] or non-coding variants impact prediction [23], where missing key binding events can distort biological insights.

The reliance on experimental data, such as ChIP-seq, for positive sequences raises additional considerations. While these datasets are generally robust, they are not immune to biases, including experimental noise or limitations in sensitivity [1,100,101]. Moreover, ChIP-seq peaks may represent indirect binding events mediated by protein complexes rather than direct TF-DNA interactions [1,102]. This complexity adds another layer of difficulty in evaluating models, highlighting the need for complementary validation strategies.

On the other hand, the inherent uncertainty in negative sequences poses challenges in developing and validating predictive models. This issue is exacerbated in highly dynamic regulatory systems, where TF binding is modulated by temporal, spatial, and environmental factors [1,103,104]. Incorporating such variability into models remains an open problem. Future approaches might leverage multi-omics data, such as ATAC-seq or single-cell transcriptomics, to refine the definition of both positive and negative sequences and improve model training as well as their evaluation.

## Data Availability

The source code to reproduce the analyses presented in this paper is available at https://github.com/InfOmics/PWM-SVM-benchmark. The datasets used in this benchmark can be accessed at https://doi.org/10.5281/zenodo.15052718. Additionally, the collection of SVM models trained on ENCODEE ChIP-seq dataset is available at https://doi.org/10.5281/zenodo.15056257.

## Key points

- Position Weight Matrices (PWMs) have been broadly employed due to their simplicity and interpretability. However, they fall short in capturing positional dependencies and intricate sequence interactions. Support Vector Machine (SVM)-based models have emerged as powerful alternatives.
- PWM-based models perform optimally with small datasets containing high-confidence sequences, especially when background data is unavailable. In contrast, SVM-based models excel in handling large datasets and long sequences, effectively capturing sequence context despite requiring extended training times.
- The inclusion of biologically relevant background data, such as DNase-seq, enhances model accuracy compared to synthetic alternatives. While SVMs offer superior predictive performance, they lack interpretability, whereas PWMs provide clear motif representations.
- A curated repository of pretrained SVM models trained on human ChIP-seq data has been developed, facilitating broader accessibility and application in regulatory genomics research.

## Supplementary file

### 1 STREME

STREME [1] is an enumerative motif discovery algorithm [2,3] specifically designed to identify Transcription Factor Binding Site (TFBS) motifs in DNA sequences. As an enumerative method, STREME focuses on detecting motifs by identifying sequences that are significantly overrepresented in a positive dataset compared to a background dataset. This is achieved by counting the approximate occurrences of all possible 4^*m*^ DNA sequences (*k*-mers), where m is the length of the motif. Then, the idea is to evaluate whether the differences in *k*-mers frequencies between the positive and background datasets are statistically significant. From the identified *k*-mers, enumerative methods construct an alignment profile, which is subsequently used to generate a corresponding Position Weight Matrix (PWM) [4,5] for motif representation. A key improvement of STREME, compared to other enumerative methods, is its computational efficiency. By leveraging Suffix Trees (STs) [6] to encode positive and negative sequences, STREME performs approximate sequence matching across thousands of sequences in a single traversal of the tree. This approach ensures that even longer motifs (up to 30 bp) can be analyzed quickly and accurately.

To evaluate statistical significance throughout the procedure, STREME employs different strategies tailored upon the characteristics of the input datasets. When primary and control sequences have identical length distributions, STREME employs the Fisher’s exact test, to evaluate whether motif sites are distributed equally between the datasets. Given the primary sequences *S*_*p*_ and the control sequences *S*_*c*_, with *s*_*p*_ ∈ *S*_*p*_ and *s*_*c*_ ∈ *S*_*c*_, the test calculates the probability that *s*_*p*_ ∈ *S*_*p*_ or more primary sequences contain a motif site, while *s*_*c*_ ∈ *S*_*c*_ or fewer control sequences contain motifs. For datasets with different length distributions, STREME utilizes the Binomial test to estimate motif significance. Suppose |*S*_*p*_| and |*S*_*c*_| are the numbers of primary and control sequences with average lengths *L*_*p*_ and *L*_*c*_, respectively. The approximate number of possible motif positions in each dataset is computed as *P*_*p*_ = *S*_*p*_(*L*_*p*_ − *m* − 1) and *P*_*p*_ = *S*_*p*_(*L*_*p*_ − *m* − 1), where *m* is the motif length. STREME then estimates the Bernoulli probability *P*_*b*_ of a motif site randomly falling in a primary sequence as 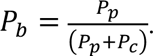 The binomial test computes the probability of observing *s* or more matches in the primary sequences, assuming a binomial distribution with success probability *P*_*b*_. The *P*-value is computed as the sum of binomial probabilities for *k* successes in trials, where *k* ranges from *s*_*p*_ to |*S*_*p*_|:

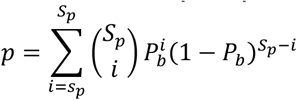

To enhance the accuracy of motif discovery, STREME further refines the search by constructing a Markov model from the control sequences. This model captures background sequence patterns, enabling the differentiation of true motifs from noise and improving the precision of the enrichment analysis.

STREME follows a multi-step, iterative process for motif discovery and Position Weight Matrix (PWM) training, beginning with the careful preparation of input sequence datasets.

In the first step, STREME processes the provided sequence datasets, which include a primary (positive) dataset and, optionally, a control (background) dataset. To ensure reproducibility, the algorithm alphabetically sorts the input sequences and then shuffles them using a fixed random seed. This approach eliminates biases introduced by sorting while maintaining consistency in results across repeated runs. If a control dataset is not supplied, STREME automatically generates a background dataset by shuffling the primary sequences. This shuffling process retains the frequencies of short *k*-mers, ensuring that the compositional characteristics of the background dataset closely resemble those of the primary dataset. This robust preparation step ensures the statistical validity of subsequent motif discovery and PWM training.

In the second step, STREME constructs a ST from a concatenated sequence of all primary and control sequences, with each sequence separated by unique delimiter characters. Each leaf of the tree corresponds to a suffix of this concatenated sequence and is labeled to indicate whether it originates from a primary or control sequence. Tree paths that do not include separator characters represent contiguous words occurring in one or more input sequences. The resulting ST allows STREME to perform rapid, scalable analysis of sequence patterns across both datasets.

In the subsequent step, STREME performs a depth-first traversal of the ST, annotating each node with the counts of primary and control sequences that contain the corresponding word. The algorithm identifies potential motifs by selecting nodes representing words within the specified width range and evaluating their statistical significance using an enrichment test to calculate *P*-values. Words with significant enrichment in the primary dataset relative to the control dataset are retained as candidate motifs. Initial seeds are then derived as valid prefixes of these selected nodes, serving as the starting point for further motif refinement.

Next, STREME refines candidate motifs by aligning good seed words into PWMs, used to evaluate other words encoded in the ST. Nodes in the suffix tree that match the PWM, or its prefixes are recorded. The statistical significance of the motif is then recalculated. This iterative refinement process continues until no further improvement in the *P*-value is observed. The most statistically significant PWM is then reported.

In the final stages, STREME evaluates the discovered motif’s ability to differentiate primary sequences from control sequences using a hold-out dataset. Statistical tests are applied to assess the motif’s classification performance, ensuring its robustness and biological relevance.

To avoid redundant detections in subsequent iterations, the identified motifs are "erased" from the input sequences by replacing their occurrences with unique separator characters. This process enables STREME to focus on uncovering additional, non-overlapping motifs in subsequent iterations, improving the diversity and comprehensiveness of the identified motif set.

### 2 MEME

MEME [7—9] is an alignment-based algorithm [2,3] designed to discover multiple motifs within unaligned genomic sequences, operating without prior assumptions. Specifically, it does not require predefined knowledge of motif width, location, sequence composition, or the number of motifs shared across input sequences. MEME focuses exclusively on contiguous motifs, allowing for mutations within the motif but prohibiting insertions or deletions. This constraint ensures that all matching subsequences have the same length, simplifying the alignment process and enabling robust motif identification.

MEME extends the Expectation Maximization (EM) algorithm to discover motifs in input sequences by introducing three novelties. (i) MEME begins motif discovery by leveraging subsequences occurring in the input sequences as starting points for motif discovery. The algorithm increases the likelihood of identifying optimal motifs compared to starting from random initializations. (ii) Unlike other motif discovery assuming one single motif occurrence per sequence, MEME allows multiple motif occurrences. This flexibility allows the algorithm to analyze sequences motifs are repeated or distributed non-uniformly. Additionally, MEME disregards sequences lacking shared motifs, enhancing its robustness against noisy data. (iii) After identifying a motif, the algorithm probabilistically erases it from data, enabling the discovery of additional distinct motifs. This iterative approach ensures that multiple motifs, even if they overlap or appear in different regions of the input sequences, can be reliably discovered.

The MEME algorithm encompasses two main steps: the Expectation (E) step and the Maximization (M) step. In the E-step, the algorithm estimates the probability of a shared motif starting at each possible position in each sequence, based on the current motif model. In the M-step, these probabilities are used to refine the motif parameters, maximizing the overall fit of the model to the data. These steps are repeated iteratively until convergence, resulting in a local maximum of the likelihood function.

However, traditional EM algorithms face several limitations. They heavily rely on initial guesses making it unclear if the true motif has been discovered, and their stopping criteria are often arbitrary. Additionally, the “one-occurrence-per-sequence” model (OOPS) [10] assumes each sequence contains exactly one motif, leading to underrepresentation of sequences with multiple motifs and overrepresentation of those without motifs. Moreover, this approach terminates after finding a single motif, limiting its scope.

MEME addresses these issues with several novelties. (i) Instead of relying on a single initial guess, MEME systematically selects starting points from all subsequences of a given length in the dataset. These subsequences are converted into letter probability matrices for EM initialization. To allow flexibility, MEME assigns a fixed probability to observed letters in the subsequences and distributes the remaining probability across other letters. This ensures the algorithm can capture motif variability. (ii) MEME assumes that sequences may contain zero, one, or multiple motif occurrences (ZOOPS model [10]), reflecting real-word biological data. However, MEME requires that the expected number of motif occurrences of the motif in the dataset is known *a priori*. MEME’s ZOOPS model reduces bias from sequences without motifs and ensures those with multiple motifs contribute appropriately. (iii) To avoid interpreting long, repeated sequences as multiple occurrences of a shorter motif, MEME enforces a constraint, where no consecutive positions can have a combined probability > 1.0. This constraint prevents overcounting artifacts. (iv) MEME optimizes computation by running a simplified EM process for each starting point, generating candidate motif models. The best candidate, based on likelihood, undergoes further refinement through a full EM process until convergence. These innovations enable MEME to discover motifs more effectively and robustly, even in complex datasets with noisy or variable sequences, making it a powerful tool for motif discovery in genomic studies.

Another key innovation of MEME is its ability to iteratively discover multiple distinct motifs in datasets containing more than one conserved motif. Unlike OOPS model, which may focus exclusively on the most conserved motif, MEME employs a probabilistic erasure mechanism to identify additional motifs.

Each position in a sequence is initially assigned a weight of 1.0, representing the probability that has not been part of a previously discovered motif. After identifying a motif, MEME updates these weights based on offset probabilities: positions with a strong match to the discovered motif have their weights significantly reduced, effectively erasing them from subsequent searches. Conversely, positions with weaker matches retain higher weights, allowing them to contribute to the discovery of additional motifs. This approach ensures that strongly conserved motifs are prioritized while still enabling weaker matches to be considered in later iterations, reducing the likelihood of missing potential motifs due to chance similarities.

In addition, MEME assesses motifs strength and significance using a log-odds matrix. This matrix facilitates the calculation of likelihood ratios to determine whether a given subsequence is more likely to originate from the motif or the background noise. Each entry in the matrix represents the log-odds score for a specific subsequence, calculated as the sum of the log-odds values for each position in the subsequence. Positive log-odds scores indicate that the subsequence is more likely generated by the motif, while negative scores suggest it belongs to the background.

By using the log-odds matrix in combination with a threshold, MEME can identify potential motif instances in unseen data and assess their similarity to previously discovered motifs, Enabling robust and reliable motif discovery and annotation.

### 3 LS-GKM

LS-GKM [11] provides an efficient approach for analyzing large-scale sequence datasets by using gapped *k*-mers to train Support Vector Machine (SVM)-based motif models. This method is particularly well-suited for capturing sequence patterns underlying transcription factor binding sites (TFBS) in a computationally efficient manner.

The central concept of LS-GKM is to differentiate between two datasets: one containing bound (positive) sequences and the other containing unbound (negative) sequences. Unlike traditional *k*-mer-based methods that rely on exact *k*-mer frequency counts, LS-GKM increases flexibility and sensitivity by incorporating gapped *k*-mers [12]. Gapped *k*-mers are subsequences that include gaps, or positions that can be skipped, allowing the model to capture patterns with subtle variations.

LS-GKM represents each sequence as a feature vector derived from the occurrences of its gapped k-mers. This feature set is defined by two parameters: the k-mer total length *l*, including gaps, and the number of informative positions in the *k*-mer *k*.

The resulting gapped *k*-mer has *l* − *k* gaps. For a sequence *s*, the feature vector is denoted as:

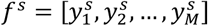

Where 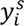 represents the number of occurrences of the *i*-th gapped *k*-mer in *s*, and *M* is the total number of possible gapped *k*-mers. This method effectively reduces the dimensionality of the feature space compared to exact *k*-mer enumeration, which would require representing all possible *k*-mer combinations. By leveraging gapped *k*-mers, LS-GKM not only captures more complex sequence patterns but also improves the model’s ability to detect motifs with subtle variations, such as those arising from sequence mutations or alternative binding preferences.

The feature vectors are subsequently used to train an SVM model with specialized kernel functions. The gapped *k*-mer (gkm) kernel is a key component, defined as the normalized inner product between the feature vectors of two sequences *s*_1_ and *s*_2_:

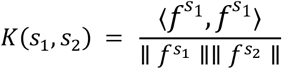

where 〈*f*^*s*_1_^, *f*^*s*_1_^ 〉 represents the inner product of the feature vectors, while ∥ *f*^*s*_1_^ ∥ and ∥ *f*^*s*_2_^ ∥ denote the *L*^2^ norms of the respective feature vectors, which ensure normalization. This kernel quantifies the similarity between sequences by evaluating the overlap of their gapped *k*-mer representations. The normalization ensures that the kernel value is not biased by the magnitude of the feature vectors, enabling fair comparison between sequences of varying lengths or *k*-mer composition.

The gkm kernel can be computed efficiently without explicitely enumerating all gapped *k*-mers, which would otherwise render direct computation intractable. The approach relies on two key observations. (i) Only the full *l*-mers present in the sequences contribute to the inner product via all gapped *k*-mers derived from them. (ii) For each pair of full *l*-mers (belonging to different sequences), only the number of mismatches between them affects their contribution to the inner product. Therefore, using these principles, the inner product of the gkm kernel can be reformulated as:

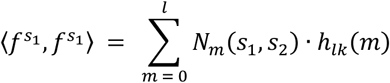

where *N*_*m*_(*s*_1_, *s*_2_) is the number of pairs of *l*-mers with *m* mismatches, referred to as the mismatch profile of *s*_1_ and *s*_2_, and *h*_*lk*_(*m*) is the contribution to the inner product for *m* mismatches, calculated as:

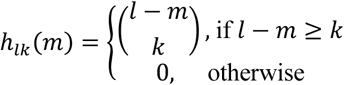

To compute the mismatch profile efficiently, a *k*-mer tree data structure is employed [13]. This approach enables the simultaneous calculation of mismatch profiles for all the sequence pairs. The k-mer tree holds all *l*-mers from all input sequences, and each leaf node represents an *l*-mer. Each leaf node stores the list of sequences containing that *l*-mer, and the frequency of the *l*-mer in each sequence. Each internal node *t*_*i*_ at depth *d* represents a subsequence of length *d*, denoted as *s*(*t*_*i*_), determined by the path from the root to *t*_*i*_. The tree is traversed using depth-first search (DFS), and during the traversal, at each node *t*_*i*_ at depth *d*, all nodes *t*_*j*_ are identified at the same depth such that *s*(*t*_*i*_) and *s*J*t*_*j*_K differ by at most *l* − *k* mismatches, and pointers to these *t*_*j*_ nodes are stored along with the number of mismatches. Moreover, at each leaf node, for every sequence *s*_*i*_ in the node’s sequence list, and for all sequences *s*_*j*_ in the pointer list, increment the corresponding mismatch profile *N*_*m*_(*s*_*i*_, *s*_*j*_) based on the number of mismatches between *s*(*t*_*i*_) and *s*(*t*_*j*_). The tree traversal speed is further optimized through multi-threading, which enables parallel processing. To achieve this, the *k*-mer tree is partitioned into smaller sub-trees, each representing a segment of the search space. These sub-trees are processed independently by separate threads, allowing for concurrent evaluation. This approach not only accelerates the computation but also ensures efficient utilization of available computational resources, significantly reducing the time required to compute the mismatch profiles for large datasets.

LS-GKM, alongside gkm kernel and classical *k*-mer count kernel, introduces four additional kernel functions [14]. The weighted gkm (wgkm) kernel leverages the observation that motifs often occur near ChIP-seq peak summits by assigning weights to *l*-mers based on their positional proximity to the summit. Consequently, the inner product of the feature vectors of two sequences *s*_1_ and *s*_2_ is computed as:

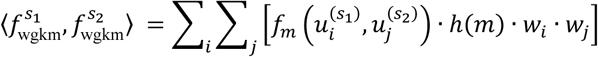

This equation extends the gkm kernel computation by incorporating weights *w*_*i*_ and *w*_*j*_, assigned to *l*-mers 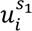 and 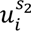 based on their positions *i* and *j* within their respective sequences. The *l*-mer pairs 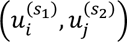 are grouped by their mismatches count *m*, with 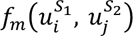 representing a function that outputs the number of *l*-mer pairs sharing *m* mismatches.

The wgkm kernel is then normalized to account for sequence lengths and scaling effects:

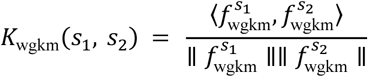

This positional weighting improves the kernel’s ability to capture biologically relevant motifs by prioritizing regions near summit peaks, enhancing predictive accuracy in motif discovery and sequence classification tasks.

The gkm radial basis (gkmrbf) kernel function builds upon the radial basis function (RBF) kernel, which is widely used to model complex non-linear relationships between feature vectors. For two generic vectors *x* and *y*, the RBF kernel is defined as:

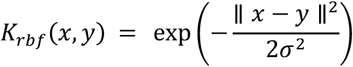

This kernel leverages the property that:

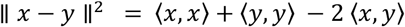

In the context of gkm-SVM, all feature vectors are normalized. Thus, for feature vectors *x* and *y* derived from sequences *s*_1_ and *s*_2_, it holds that:

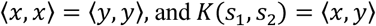

Substituting these values and defining 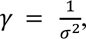 the gkmrbf kernel for sequences *s*_1_ and *s*_2_ is formulated as:

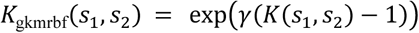

The final kernel introduced by LS-GKM is the weighted gkmrbf (wgkmrbf) kernel, which combines the principles of the gkmrbf and weighted gkm kernels. It is computed as:

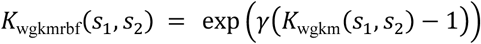

By incorporating the weighting mechanism of the wgkm kernel with the non-linear modeling capabilities of the RBF kernel, the wgkmrbf kernel provides a powerful tool for capturing complex sequence dependencies while emphasizing positional relevance. This makes it particularly suitable for tasks where motif positional information plays a crucial role.

### 4 Sequence Datasets selection and Processing

In this study, we utilized optimal IDR-thresholded ChIP-seq peaks from the ENCODE database [15] (snapshot from November 2023) to analyze 59 distinct transcription factors (TFs) representing different DNA-binding domains [16]. The ChIP-seq data were provided in the narrowPeak format, an ENCODE-specific BED-like file format commonly used for TFs due to the typically narrow regions they bind.

To extract sequence data in FASTA format from the input BED files, we used the *getfasta* command from BEDTools [17]. This command retrieves the sequences defined by the BED interval coordinates and creates corresponding FASTA entries in the output file, using the reference genome provided as input.

To obtain real biological background data, we downloaded DNase-seq peaks in BED format from ENCODE [18]. Overlaps between DNase-seq features and ChIP-seq peaks were eliminated using BEDTools’ intersect and subtract functionalities. The subtract command removed the entire DNase-seq feature if any overlap with ChIP-seq regions was detected, ensuring a clean set of DNase-seq peaks for background sequences.

For synthetic background datasets, we employed the fasta-shuffle-letters tool from the MEME Suite [19], which generates shuffled versions of input positive FASTA sequences while preserving dinucleotide frequencies. This tool uses the uShuffle algorithm [20], a sequence analysis method that ensures uniform random permutations of biological sequences, maintaining exact k-mer counts. The uShuffle algorithm constructs the de Bruijn graph of order *k* for the input sequence and generates a Eulerian path through this graph using Wilson’s algorithm [21,22]. The resulting output is a randomized version of the original sequence that retains *k*-mer properties.

Both the biological and synthetic background generation procedures were applied consistently to prepare train and test datasets for the study.

### 5 Training Position Weight Matrices and SVM-based Models

ChIP-seq data was obtained from the ENCODE database [15] and refined by excluding sequences that did not belong to canonical chromosomes. To ensure complete independence between training and testing datasets, sequences from chromosome 2 were reserved exclusively for testing, while those from all other chromosomes were used solely for training both PWM- and SVM-based models. The entire analysis was conducted with default parameters, which were explicitly documented to ensure full reproducibility of the results.

For training position weight matrices (PWMs), MEME was executed with its default parameters:

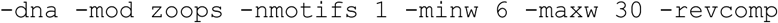

This configuration identified a single DNA motif per sequence, with a width ranging from 6 to 30 bp, accounting for reverse complements and allowing for zero or one occurrence of the motif per sequence. STREME, on the other hand, was run with the following default settings:

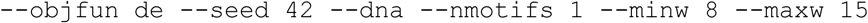

This setup aimed to identify discriminative motifs that differentiate positive from negative sequences, with strict length constraints. The --objfun de (Differential Enrichment) parameter scored motifs based on the enrichment of their occurrences in primary sequences compared to control sequences.

For training SVM-based models, the gkmtrain function from LS-GKM was employed with default parameters. This included the use of weighted gkm kernel (**Supplementary File Section 3**) with a word length of 11 and allowing up to 3 mismatches.

The MEME and STREME pipelines produced PWMs, capturing the probabilistic structure of identified motifs. Meanwhile, LS-GKM generated an SVM model file along with metadata, enabling the classification of genomic regions based on their sequence features.

### 6 Scoring Sequences

To evaluate the predictive performance of the models and identify motif occurrences, we applied distinct scoring methods tailored to SVM-based models (LS-GKM) and PWM-based models (MEME and STREME).

For PWM models, we utilized FIMO [23], which scans input sequences against a PWM. FIMO identifies motif occurrences by calculating the log-likelihood ratio (LLR), which quantifies how well a subsequence aligns with the PWM compared to a background nucleotide distribution. The LLR is then converted into a *P*-value, reflecting the statistical significance of the match. FIMO pipeline involves three main steps. (i) FIMO compares subsequences against the PWM to evaluate their alignment. (ii) Then the algorithm quantifies the alignment strength relative to background expectations calculating LLRs. (iii) Finally, FIMO converts LLRs into probabilities, emphasizing statistically significant motif occurrences. The highest score for each sequence indicated the strength and significance of its alignment with the PWM, with scores interpreted in the context of statistical probabilities. This ensured that only biologically meaningful motifs were highlighted, minimizing false positives. We applied FIMO with default settings and configured --max-stored-scores to enable high-throughput analysis without compromising precision.

For SVM-based models, we used LS-GKM’s gkmpredict [11] function to score both positive (ChIP-seq peaks) and negative (background) sequences. Using the SVM model trained via gkmtrain, gkmpredict evaluates each sequence with the features learned during training. Higher scores reflected stronger resemblance to the trained motif model. These scores were computed by leveraging the intrinsic kernel function of the LS-GKM framework, effectively capturing the gapped *k*-mer patterns characteristic of transcription factor binding sites.

### 7 Analyzing Performance of SVM-based Motif Models Collection

The SVM models trained on the comprehensive collection of human transcription factor ChIP-seq data from ENCODE [15] (Section 6.2) demonstrated robust performance. Across 2,393 ENCODE experiments performed on different cell lines and tissues, the models achieved mean scores of 0.77 for AUPRC, 0.96 for AUROC, and

0.69 for F1-scores. Certain transcription factors consistently performed well across multiple metrics, with ZNF426 emerging as a top performer in AUPRC, AUROC, and F1. Similarly, ZNF274 and ZNF433 ranked among the highest for AUPRC and F1. These factors belong to the C2H2 zinc finger protein family, which binds specifically to long, unique DNA recognition sequences [24], likely contributing to their superior performance.

These findings are consistent with observations from LS-GKM k-mer weighting for CTCF [25] (Supplementary Figure 8), where the model assigns greater weight to regions near the peak summit in positive sequences, suggesting the presence of a long, distinct TFBS. Similarly, transcription factors such as GATA1 [26] (Supplementary Figure 9), GATA2 [27] (Supplementary Figure 10), and JUND [28] (Supplementary Figure 11) exhibit *k*-mer weighting patterns in regions surrounding the peak summit, possibly indicating the presence of cis-regulatory elements that may facilitate TF–DNA binding [29, 30], supporting a more complex binding grammar. Conversely, the poorest-performing TFs, such as p53 [31] (Supplementary Figure 12), exhibit a random pattern in weight assignment, further emphasizing the relationship between LS-GKM learned weights and TFBS detection capabilities.An analysis of sequence length, peak count and total experiment size (measured in base pairs) revealed distinct performance trends (Supplementary Figure 13). A strong positive correlation was observed between sequence length and all performance metrics, suggesting that longer sequences enhance predictive accuracy. Notably, while AUROC remained consistently high, AUPRC and F1 showed greater improvements with increasing sequence length, reflecting gains in precision and recall.

In contrast, the relationship between peak count and performance was more complex. LS-GKM performance improved with a rising number of sequences, displaying an initial increase that plateaued as peak counts grew further. Correlation analysis highlighted this difference, showing a stronger association between sequence length and performance metrics (AUPRC: 0.44, AUROC: 0.43, F1: 0.54) than between peak count and these metrics (AUPRC: 0.20, AUROC: 0.19, F1: 0.06). To provide a comprehensive overview, the total experiment size also exhibits a weak correlation with the previously mentioned metrics (AUPRC: 0.30, AUROC: 0.26, F1: 0.20).

The observed trends underscore the importance of sequence characteristics in model performance. Longer sequences likely provide more comprehensive and informative patterns for SVM models to learn, enhancing predictive accuracy. The relatively modest impact of peak count may reflect diminishing returns as data redundancy increases. These findings emphasize the need to optimize sequence selection, balancing quantity and quality to maximize model performance. Furthermore, the exceptional performance of C2H2 zinc finger proteins aligns with their well-documented binding specificity, illustrating how biological features can influence computational outcomes. Future studies could explore whether these trends hold across other transcription factor families and datasets, guiding further refinements in model training and data preprocessing strategies.

---

**Supplementary Figure 1.**
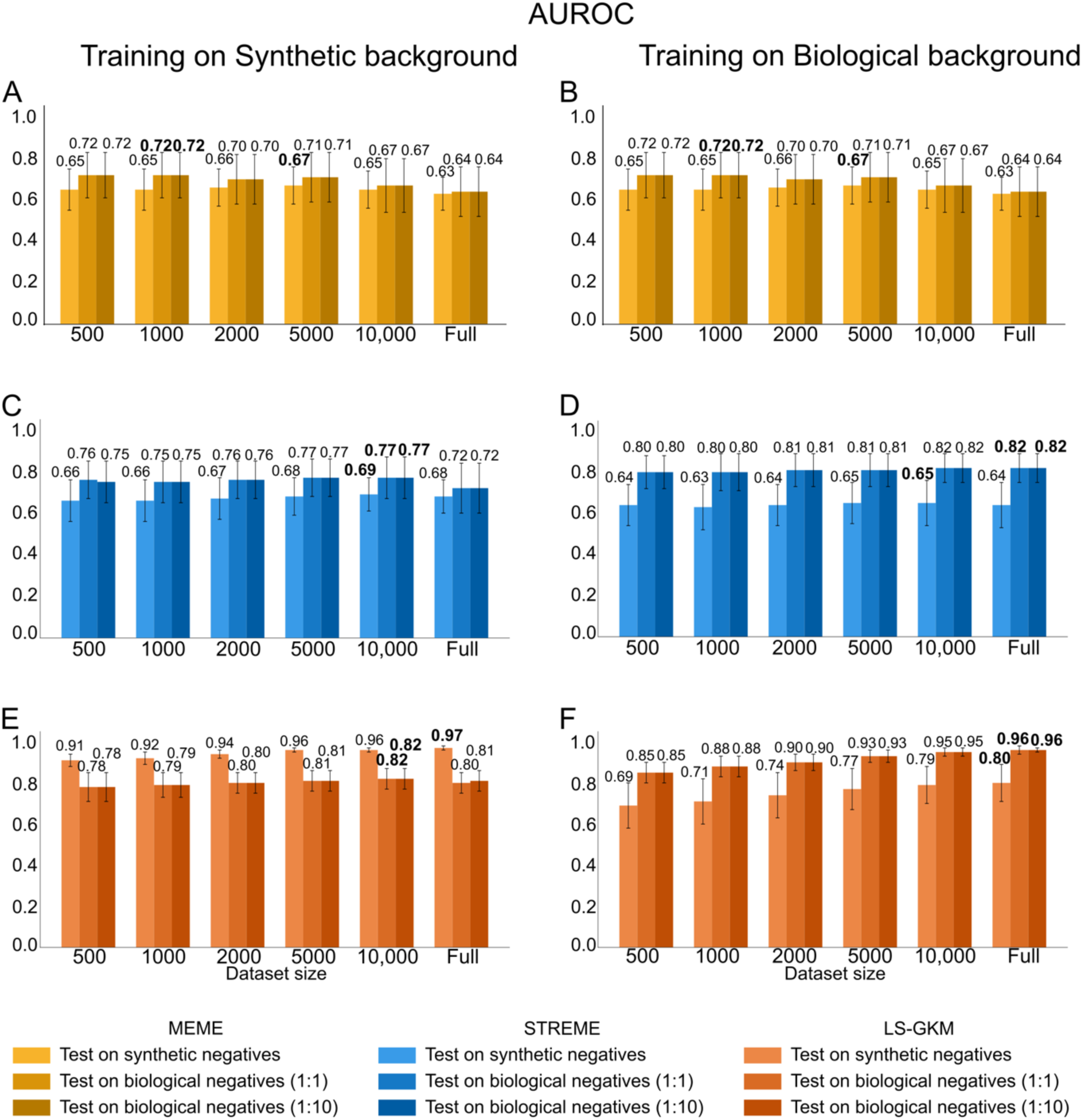
Average AUROC values from models trained across 59 datasets with varying background and dataset sizes. **(A, B)** AUROC performance of MEME’s PWMs trained on synthetic **(A)** or real biological **(B)** background data, tested on synthetic negative sequences and real biological negatives (1:1 and 1:10 positive-negative ratios). **(C, D)** AUROC performance of STREME’s PWMs trained on synthetic **(C)** or real biological **(D)** background data, evaluated under the same testing conditions. **(E, F)** AUROC performance of SVM-based models trained on synthetic **(E)** or real biological **(F)** background data, tested similarly.

**Supplementary Figure 2.**
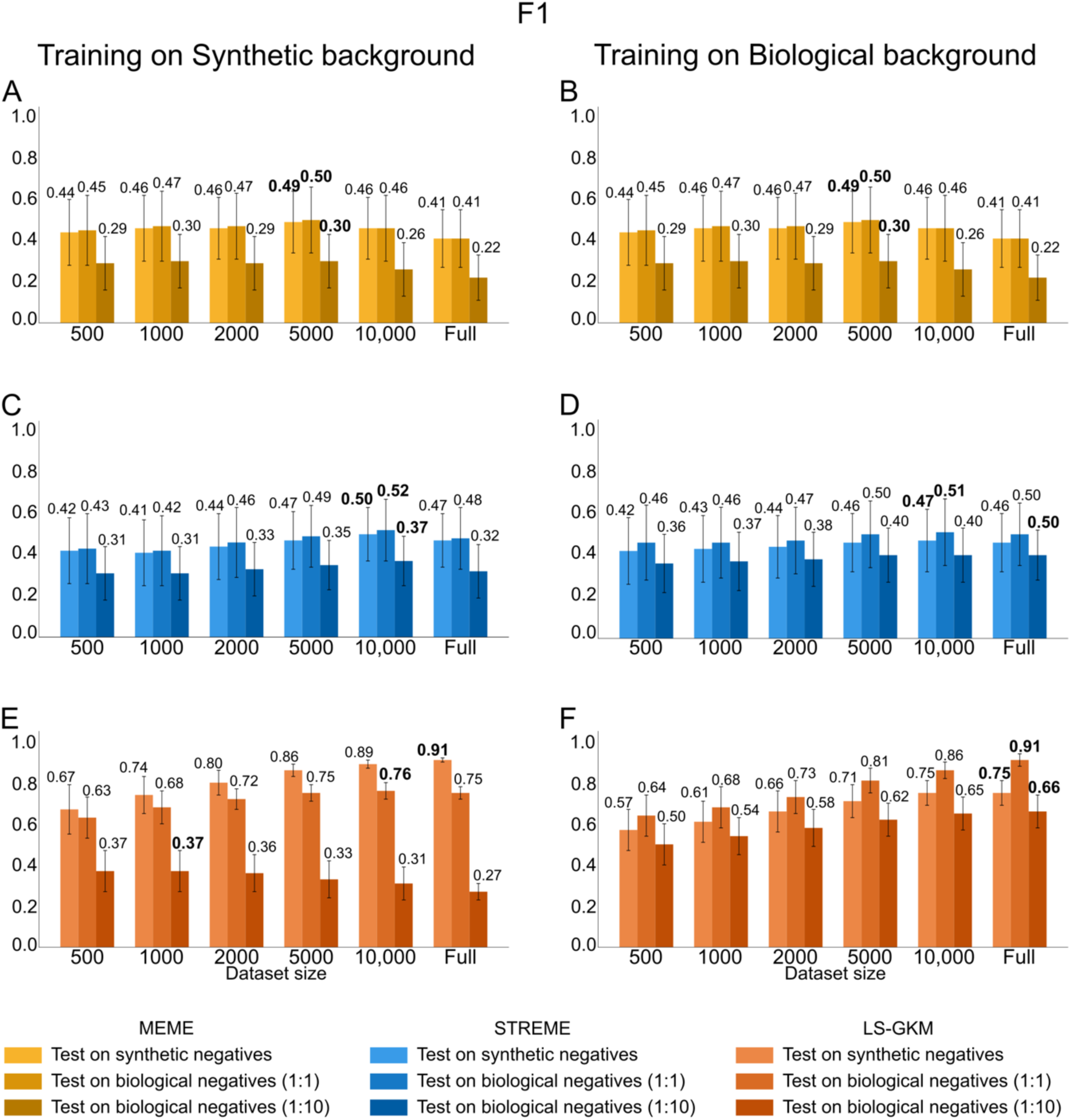
Average F1-score values from models trained across 59 datasets with varying background and dataset sizes. **(A, B)** F1-score performance of MEME’s PWMs trained on synthetic **(A)** or real biological **(B)** background data, tested on synthetic negative sequences and real biological negatives (1:1 and 1:10 positive-negative ratios). **(C, D)** F1-score performance of STREME’s PWMs trained on synthetic **(C)** or real biological **(D)** background data, evaluated under the same testing conditions. **(E, F)** F1-score performance of SVM-based models trained on synthetic **(E)** or real biological **(F)** background data, tested similarly.

**Supplementary Figure 3.**
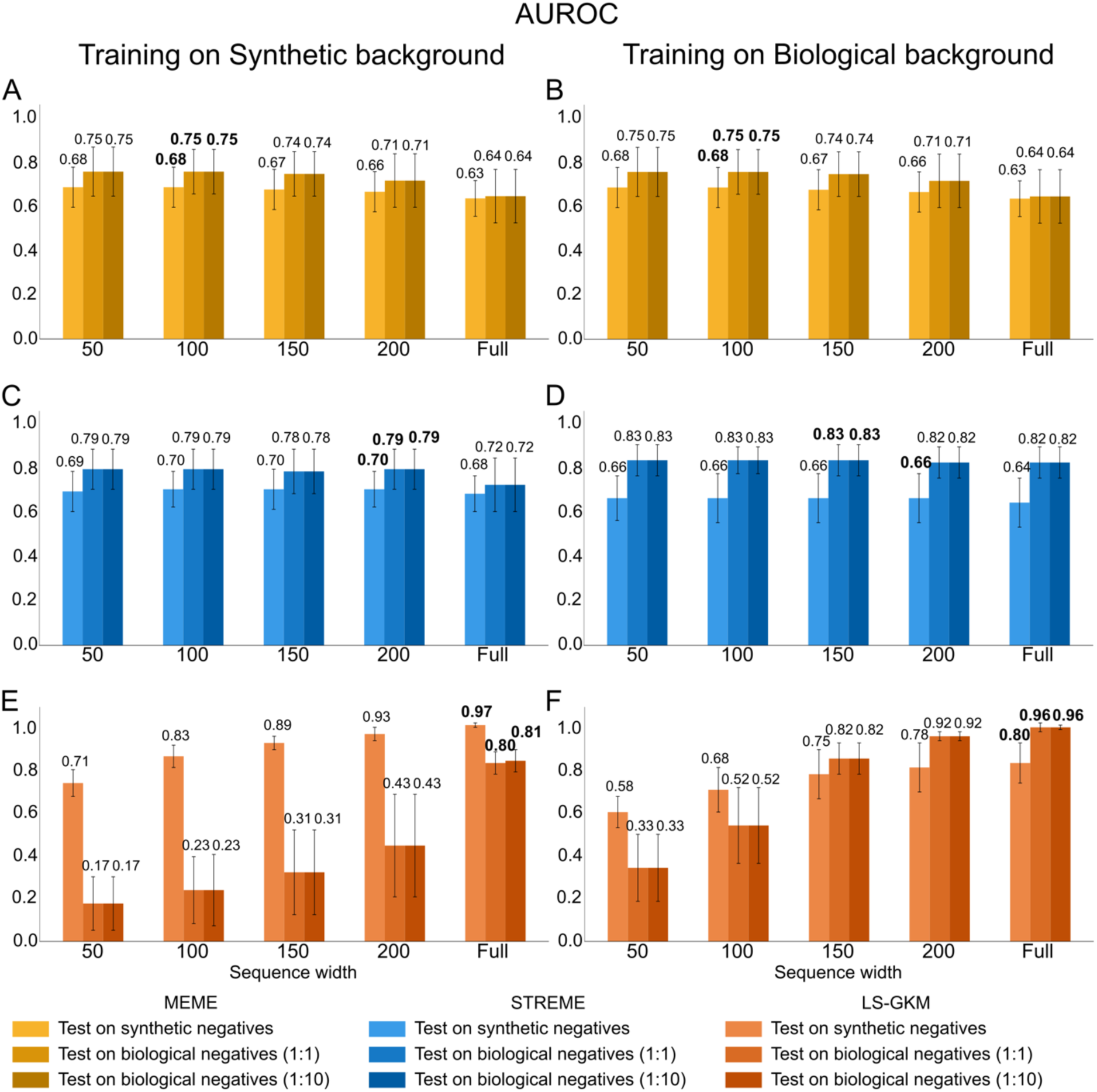
Average AUROC values from models trained across 59 datasets with varying background and sequence widths. **(A, B)** AUROC performance of MEME’s PWMs trained on synthetic **(A)** or real biological **(B)** background data, tested on synthetic negative sequences and real biological negatives (1:1 and 1:10 positive-negative ratios). **(C, D)** AUROC performance of STREME’s PWMs trained on synthetic **(C)** or real biological **(D)** background data, evaluated under the same testing conditions. **(E, F)** AUROC performance of SVM-based models trained on synthetic **(E)** or real biological **(F)** background data, tested similarly.

**Supplementary Figure 4.**
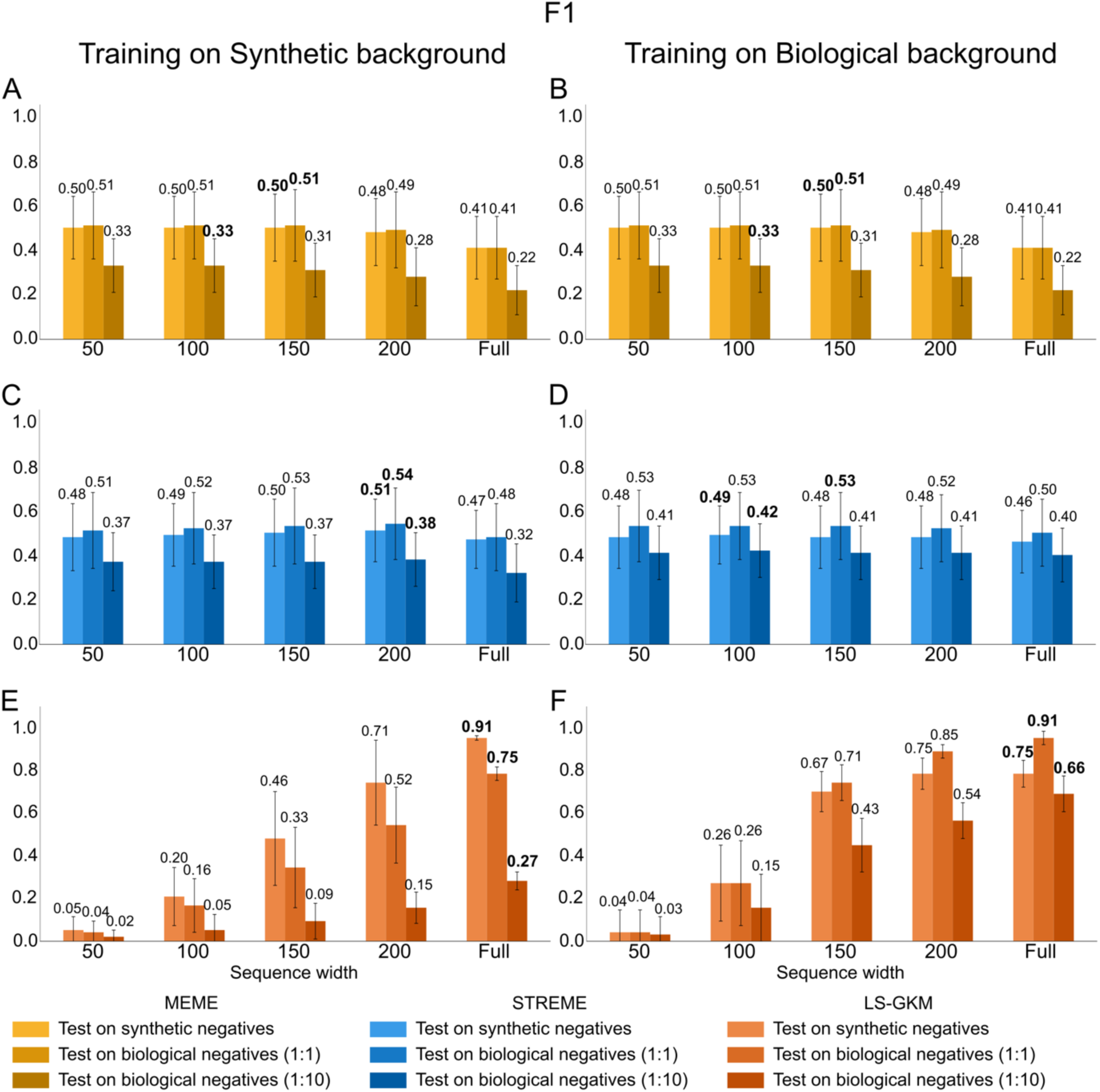
Average F1-score values from models trained across 59 datasets with varying background and sequence widths. **(A, B)** F1-score performance of MEME’s PWMs trained on synthetic **(A)** or real biological **(B)** background data, tested on synthetic negative sequences and real biological negatives (1:1 and 1:10 positive-negative ratios). **(C, D)** F1-score performance of STREME’s PWMs trained on synthetic **(C)** or real biological **(D)** background data, evaluated under the same testing conditions. **(E, F)** F1-score performance of SVM-based models trained on synthetic **(E)** or real biological **(F)** background data, tested similarly.

**Supplementary Figure 5.**
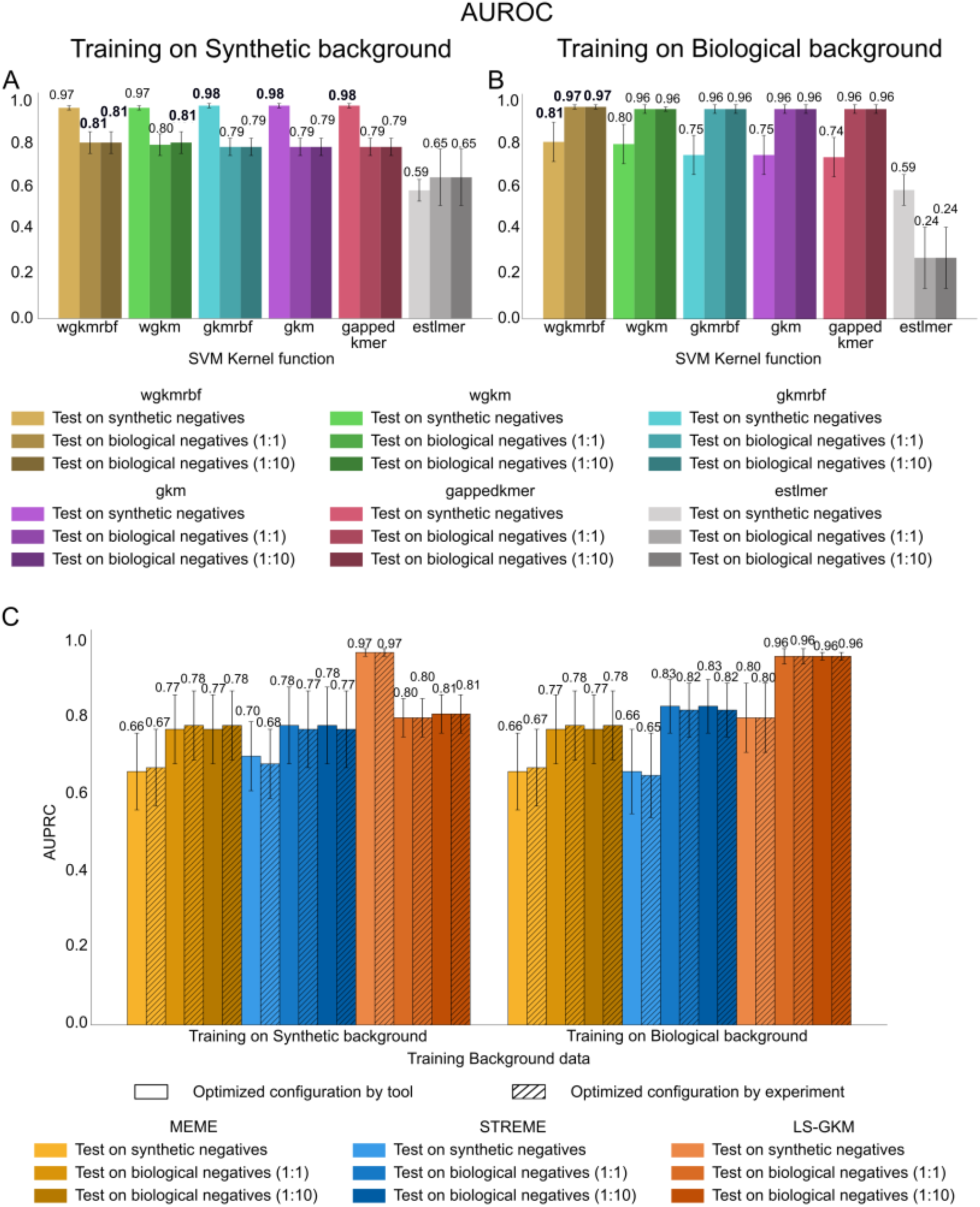
Average AUROC values for SVM-based models trained across 59 datasets, comparing different LS-GKM kernel functions and optimized training configurations. **(A, B)** Performance of kernel functions available in LS-GKM when trained on synthetic **(A)** and real biological **(B)** background sequences. **(C)** Comparison of model performance using two optimization strategies: a globally optimized tool-specific configuration versus per-experiment fine-tuning.

**Supplementary Figure 6.**
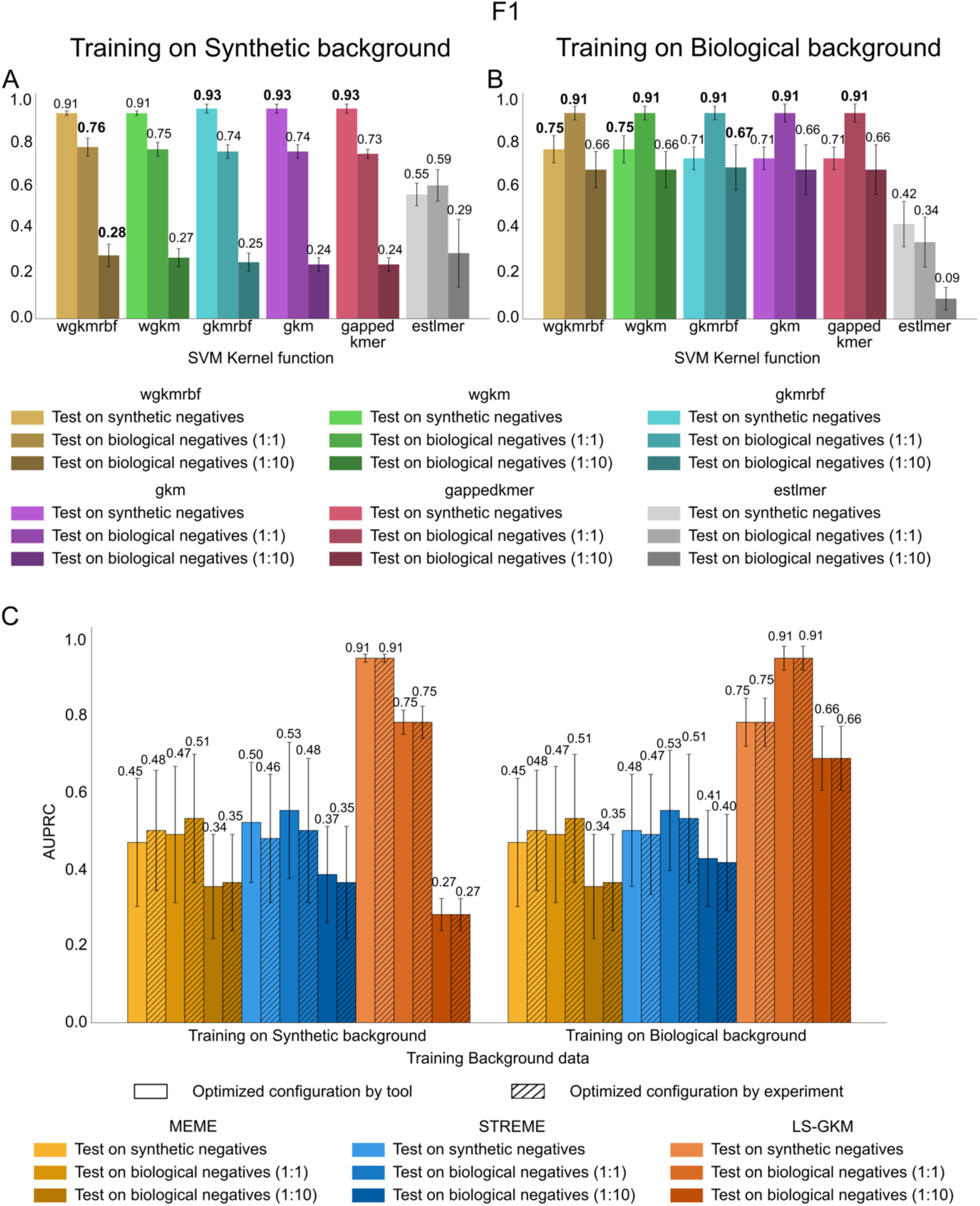
Average F1-score values for SVM-based models trained across 59 datasets, comparing different LS-GKM kernel functions and optimized training configurations. **(A, B)** Performance of kernel functions available in LS-GKM when trained on synthetic **(A)** and real biological **(B)** background sequences. **(C)** Comparison of model performance using two optimization strategies: a globally optimized tool-specific configuration versus per-experiment fine-tuning.

**Supplementary Figure 7.**
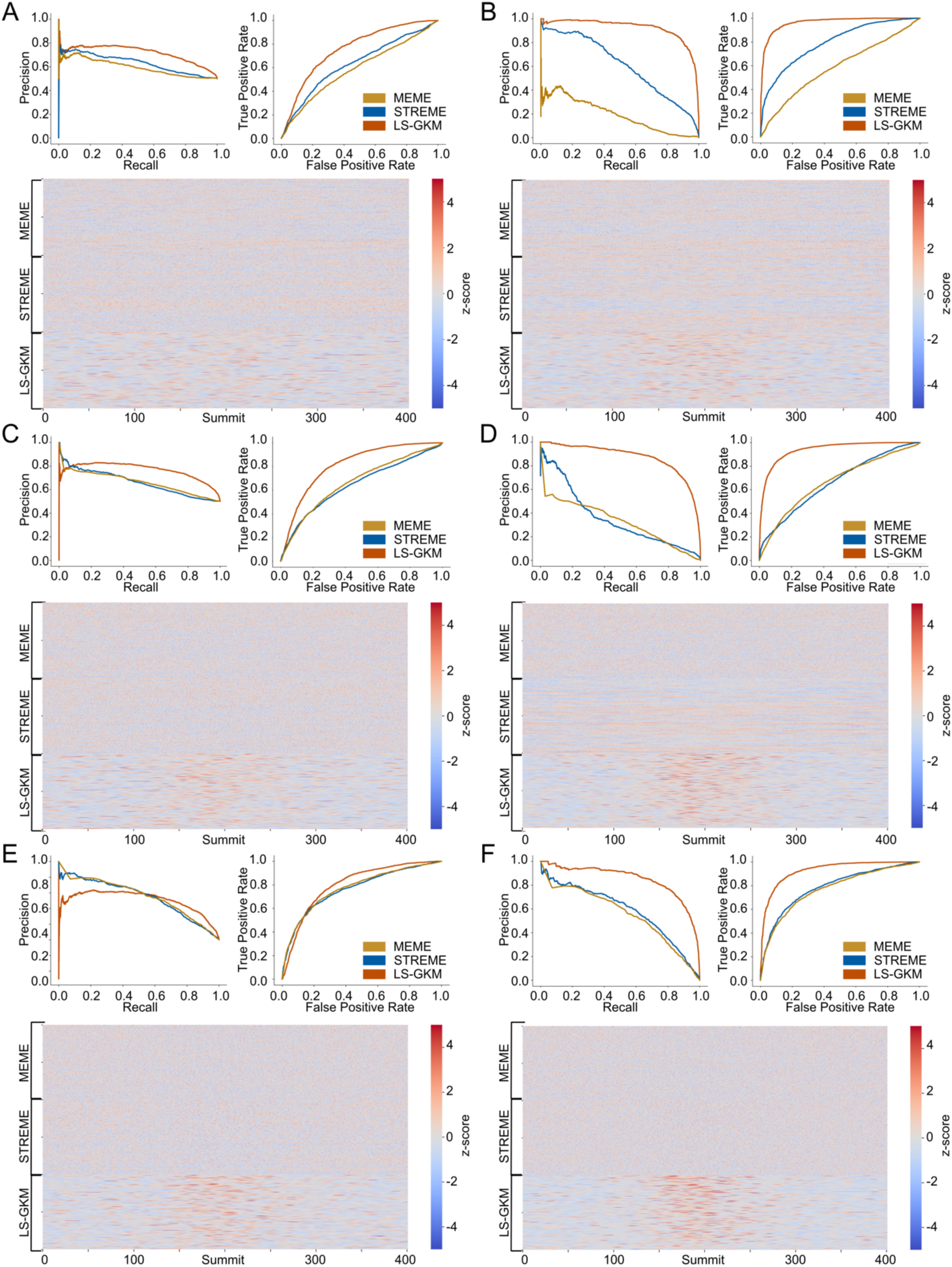
Comparison of transcription factor binding predictions by MEME, STREME, and LS-GKM trained on synthetic (left) and real biological (right) background sequences. The panels present results for the worst-performing TFs of each tool: **(A, B**) MYBL2 for MEME, (**C, D**) SOX13 for STREME, and (**E, F**) TEAD4 for LS-GKM. Heatmaps illustrate the positional distribution of motif scores across test sequences, highlighting motif enrichment patterns near peak summits. ROC (top left) and precision-recall (top right) curves summarize predictive performance

**Supplementary Figure 8.**
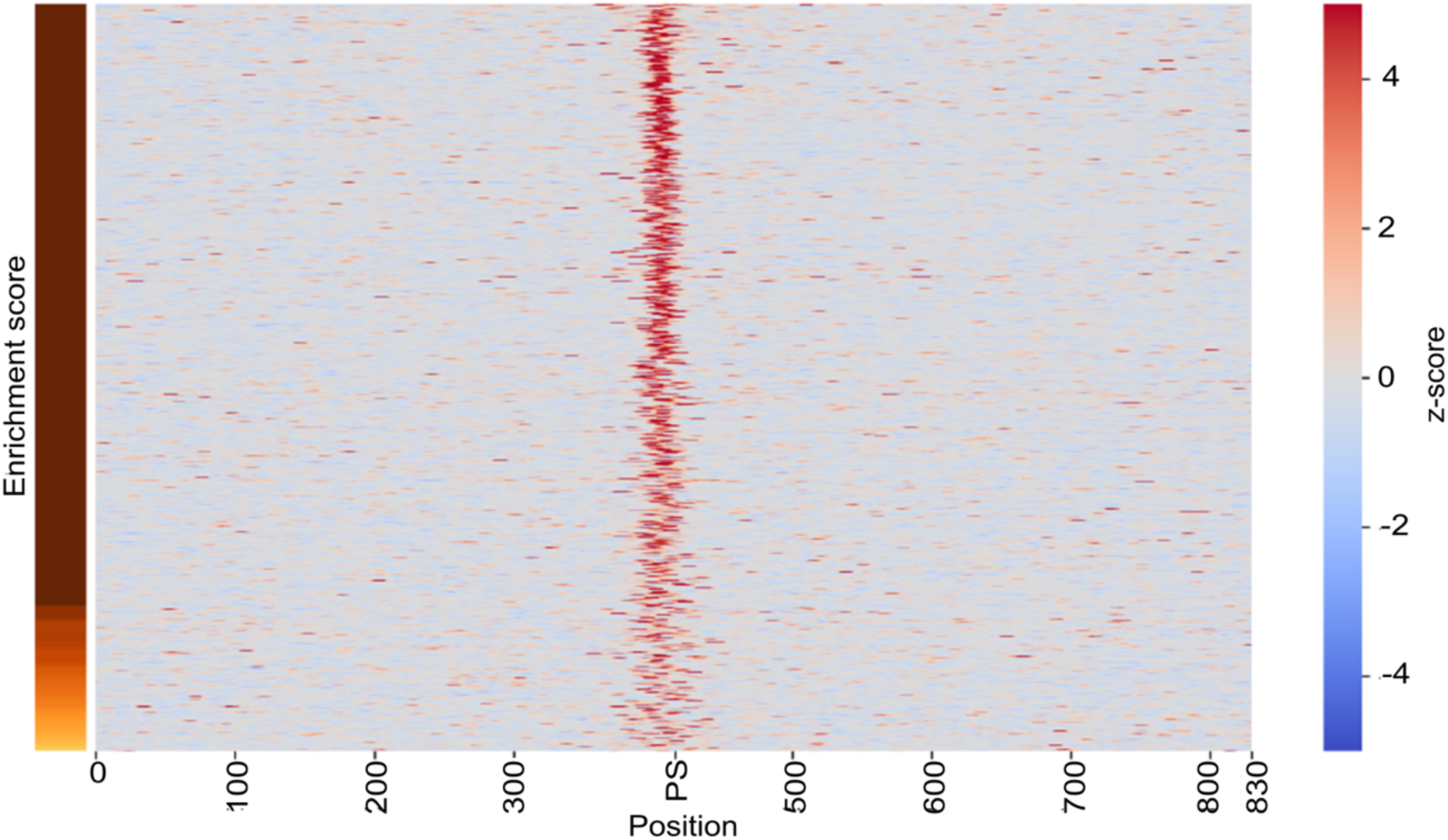
LS-GKM *k*-mer weighting for CTCF. Color scale on the right represents the z-score of the weights, while the normalized enrichment score is shown on the left.

**Supplementary Figure 9.**
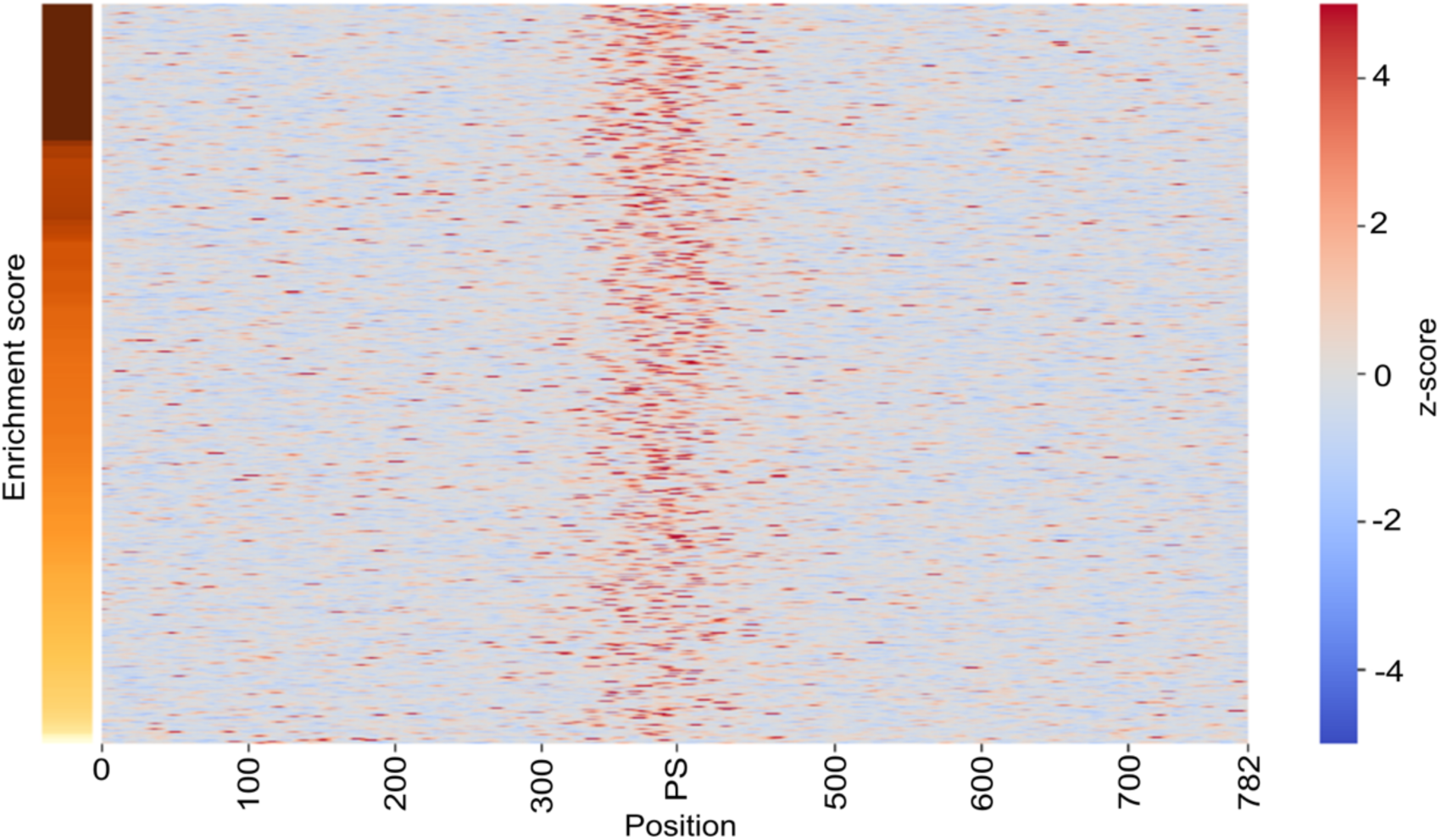
LS-GKM *k*-mer weighting for GATA1. Color scale on the right represents the z-score of the weights, while the normalized enrichment score is shown on the left.

**Supplementary Figure 10.**
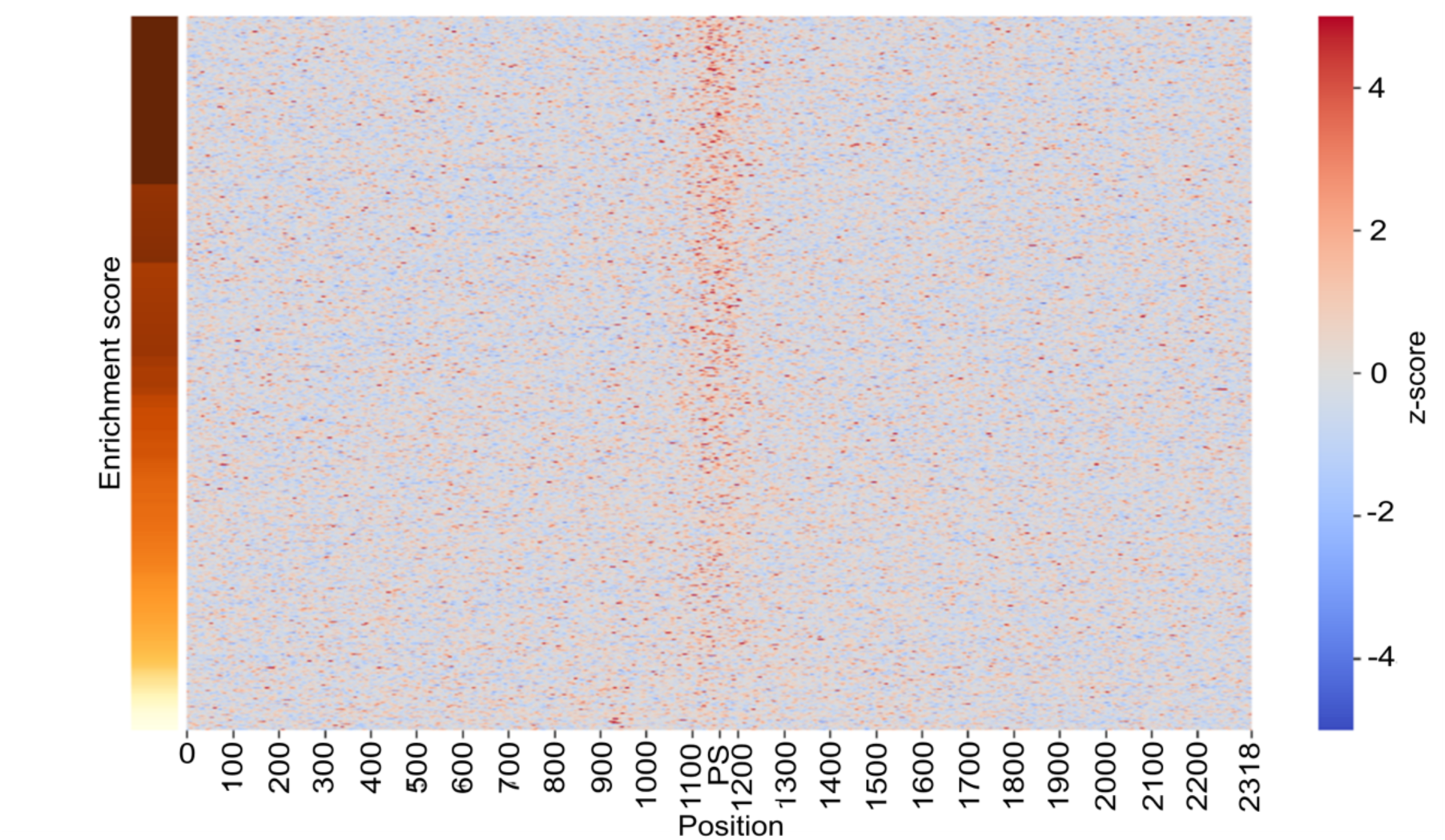
LS-GKM *k*-mer weighting for GATA2. Color scale on the right represents the z-score of the weights, while the normalized enrichment score is shown on the left.

**Supplementary Figure 11.**
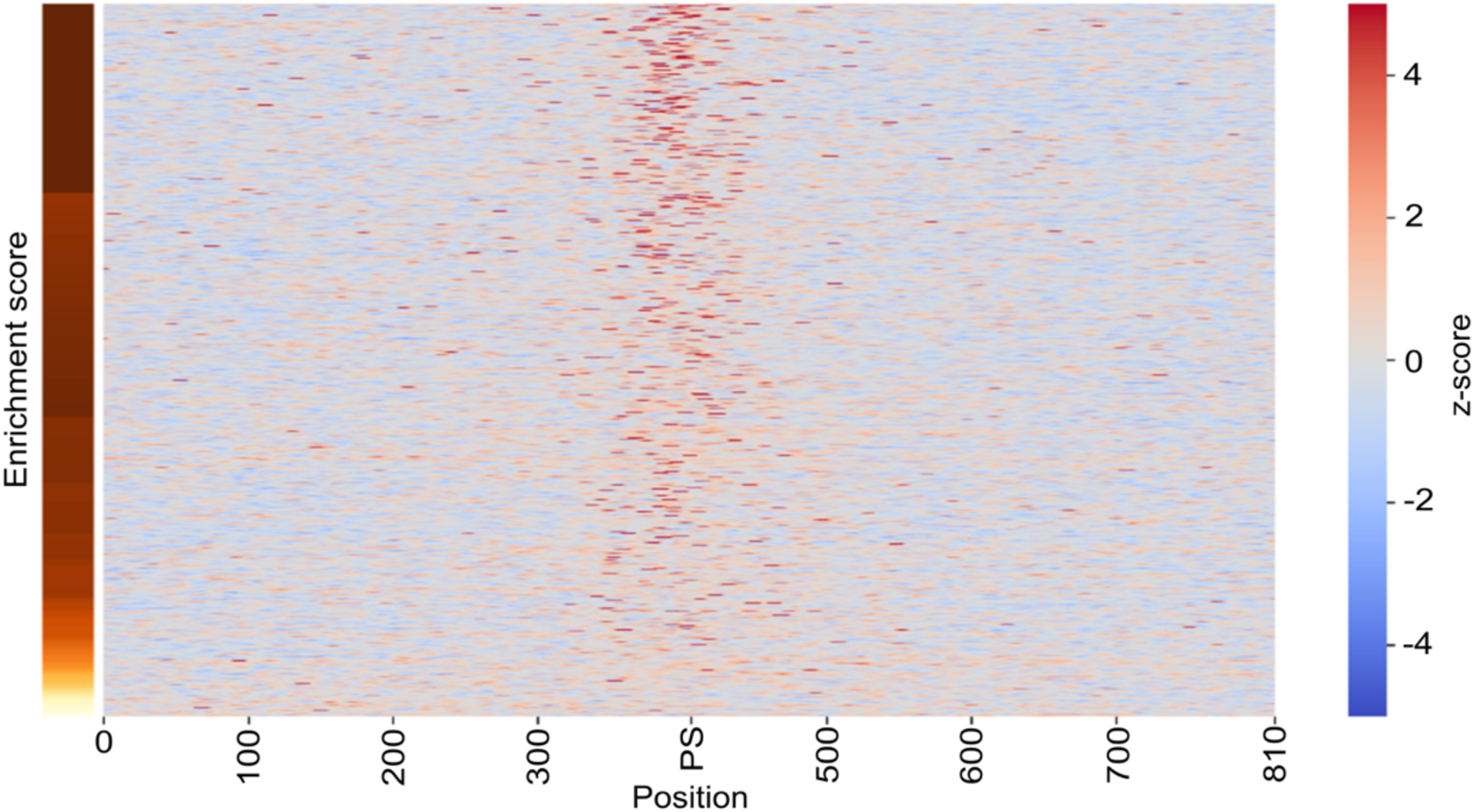
LS-GKM *k*-mer weighting for JUND. Color scale on the right represents the z-score of the weights, while the normalized enrichment score is shown on the left.

**Supplementary Figure 12.**
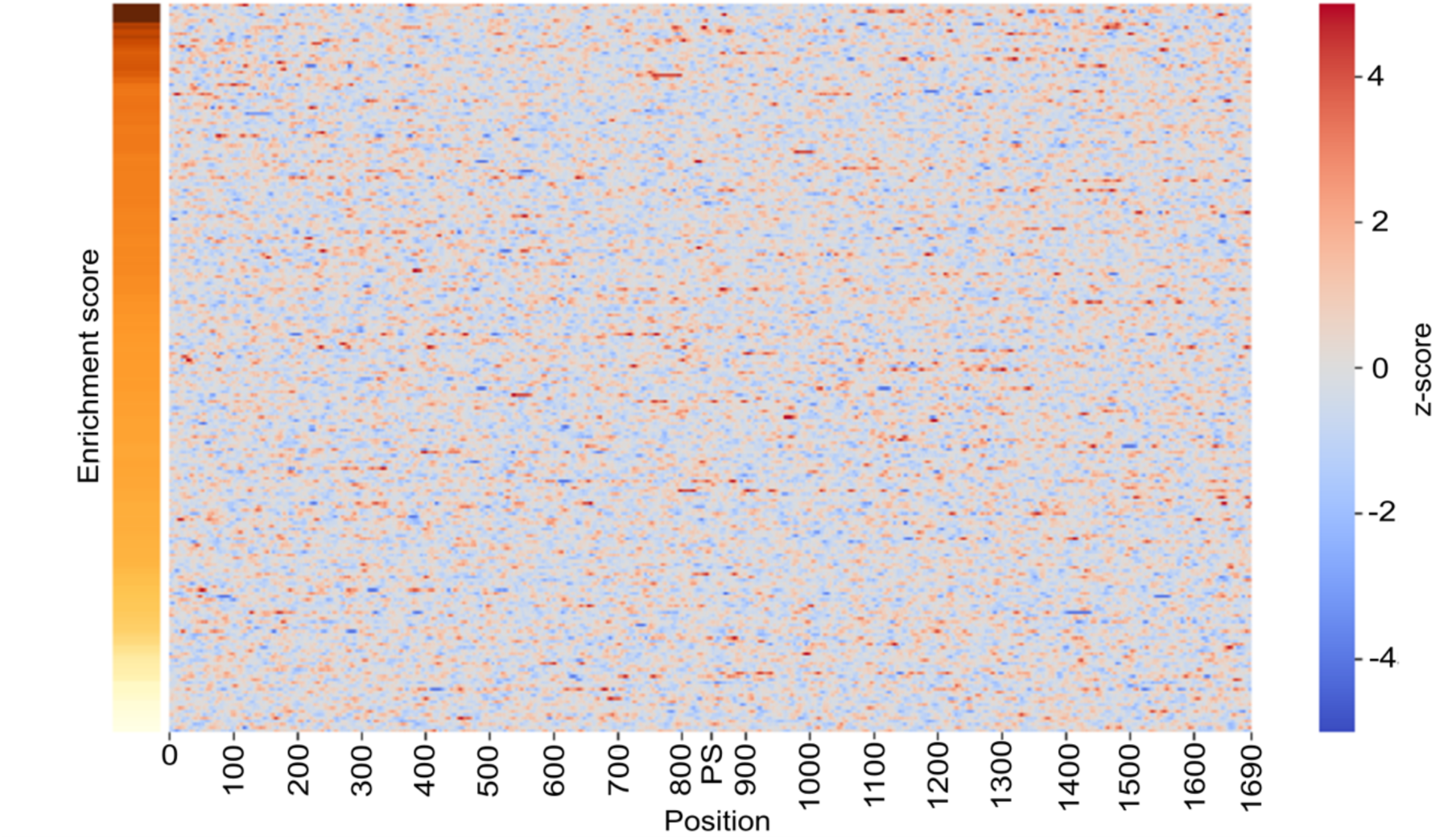
LS-GKM *k*-mer weighting for p53. Color scale on the right represents the z-score of the weights, while the normalized enrichment score is shown on the left.

**Supplementary Figure 13.**
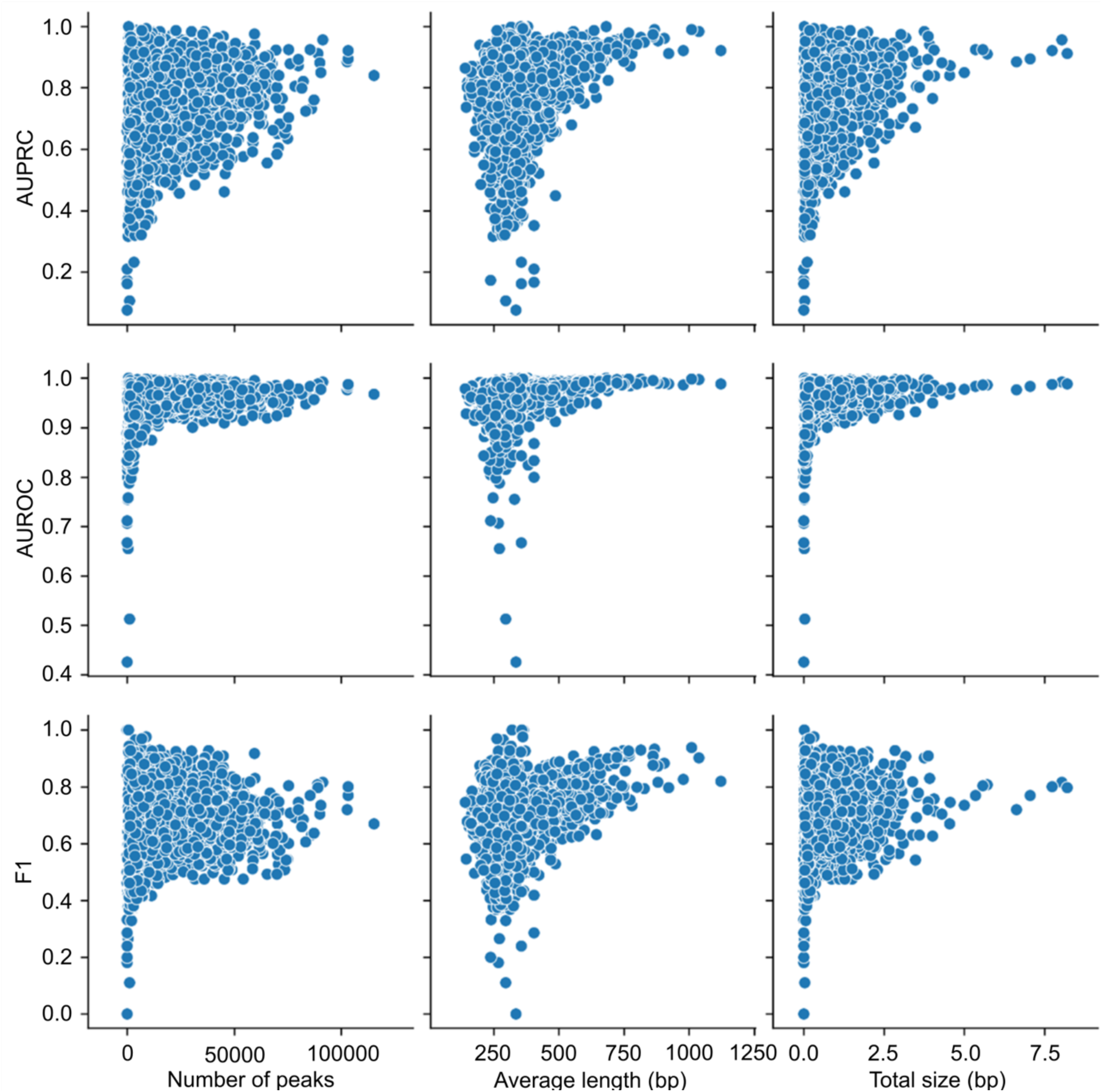
Correlation between number of peaks, average length (bp), total size (bp) and AUPRC, AUROC and F1.

**Supplementary Table 1.**
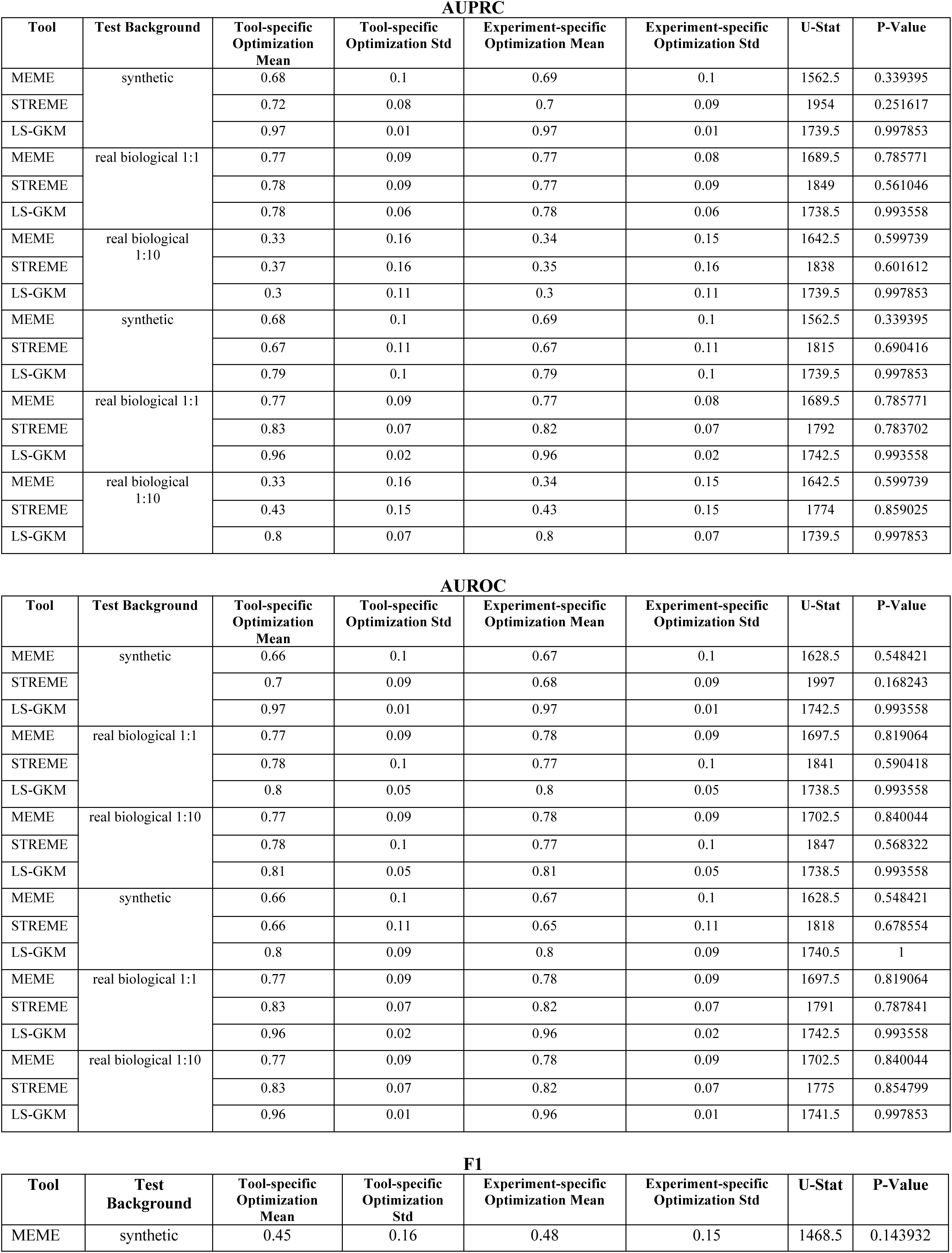
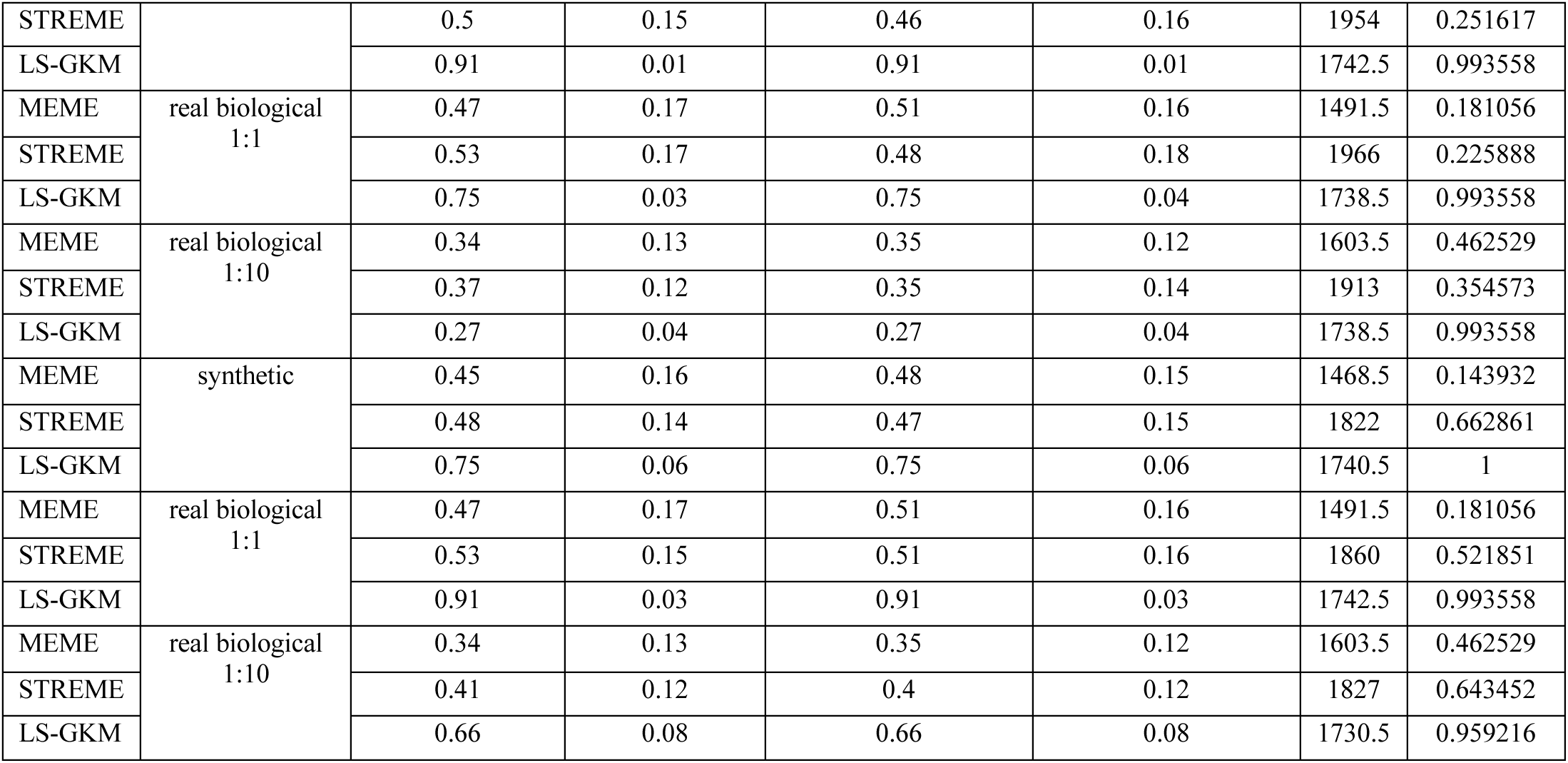
Comparison of Tool-Specific Optimization vs. Per-Experiment Optimization. This table provides general statistics on the Tool-Specific Optimization and Per-Experiment Optimization results for each training scenario.

**Supplementary Table 2.**
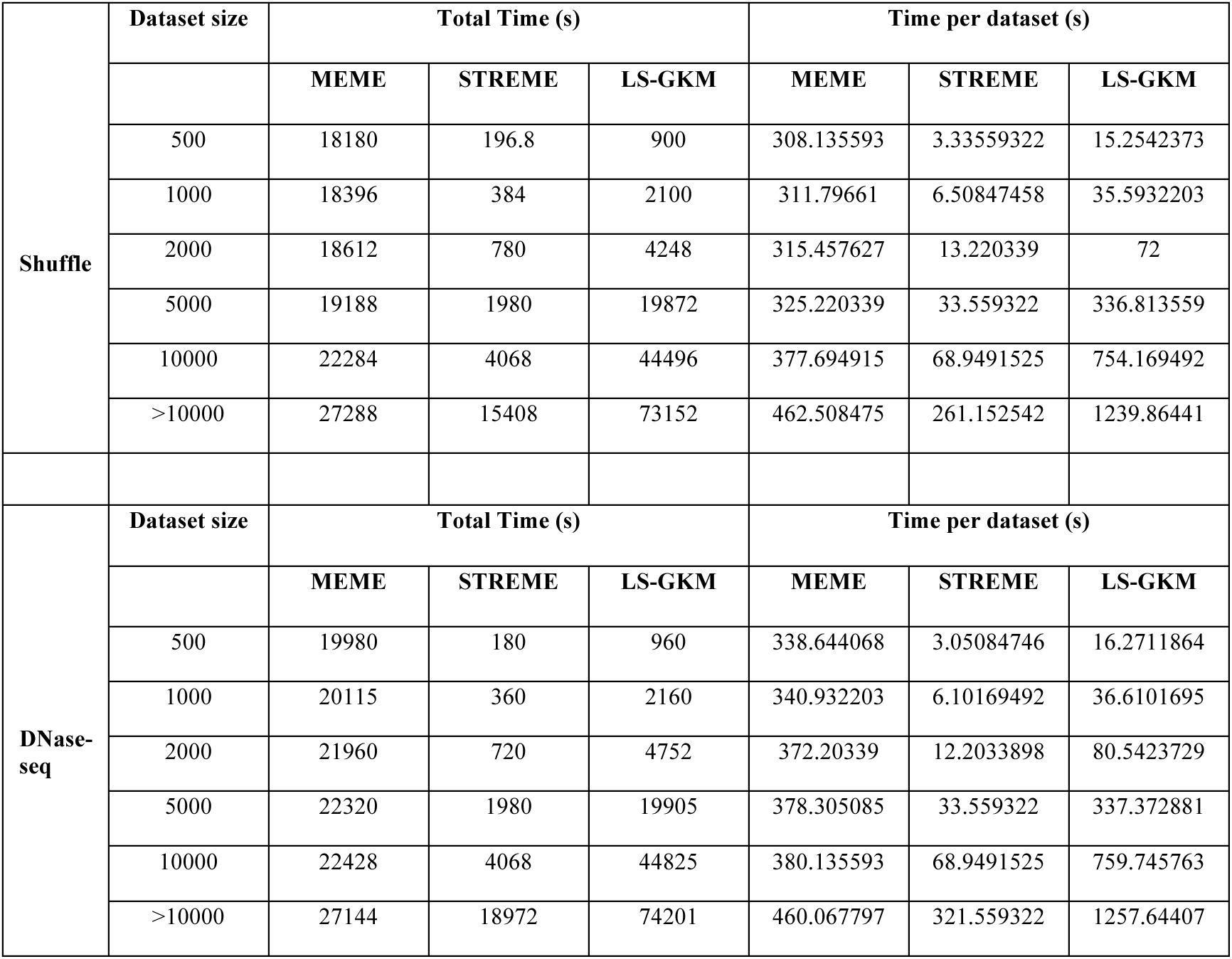
Comparison of Running Time Across Tools Based on Dataset Size. This table presents a runtime analysis for each tool relative to varying dataset sizes.

**Supplementary Table 3.** Comparison of Running Time Across Tools Based on Sequence Width. This table presents a runtime analysis for each tool relative to varying sequence widths.

|  | Sequence width | Total Time (s) |  |  | Time per dataset (s) |  |  |
| --- | --- | --- | --- | --- | --- | --- | --- |
|  |  | MEME | STREME | LS-GKM | MEME | STREME | LS-GKM |
| Shuffle | 50 | 12672 | 7884 | 66312 | 214.779661 | 133.627119 | 1123.9322 |
|  | 100 | 29088 | 16020 | 68436 | 493.016949 | 271.525424 | 1159.9322 |
|  | 150 | 29448 | 12312 | 70236 | 499.118644 | 208.677966 | 1190.44068 |
|  | 200 | 26136 | 15948 | 73080 | 442.983051 | 270.305085 | 1238.64407 |
|  | > 200 | 27288 | 19656 | 77400 | 462.508475 | 333.152542 | 1311.86441 |
|  | 50 | 12672 | 7884 | 66312 | 214.779661 | 133.627119 | 1123.9322 |
| DNase-seq | Sequence width | Total Time (s) |  |  | Time per dataset (s) |  |  |
|  |  | MEME | STREME | LS-GKM | MEME | STREME | LS-GKM |
|  | 50 | 12888 | 12528 | 66348 | 218.440678 | 212.338983 | 1124.54237 |
|  | 100 | 29988 | 22212 | 69264 | 508.271186 | 376.474576 | 1173.9661 |
|  | 150 | 26424 | 26748 | 70524 | 447.864407 | 453.355932 | 1195.32203 |
|  | 200 | 25884 | 21672 | 76032 | 438.711864 | 367.322034 | 1288.67797 |
|  | > 200 | 27072 | 33444 | 81108 | 458.847458 | 566.847458 | 1374.71186 |
|  | 50 | 12888 | 12528 | 66348 | 218.440678 | 212.338983 | 1124.54237 |

## References

1. Lambert SA, Jolma A, Campitelli LF, et al. The Human Transcription Factors. Cell 2018; 172:650–65.

2. Reimold AM, Iwakoshi NN, Manis J, et al. Plasma cell differentiation requires the transcription factor XBP-1. Nature 2001; 412:300–7.

3. Lee TI, Young RA. Transcriptional Regulation and Its Misregulation in Disease. Cell 2013; 152:1237–51.

4. Whitfield TW, Wang J, Collins PJ, et al. Functional analysis of transcription factor binding sites in human promoters. Genome Biology 2012; 13:R50.

5. Gotea V, Visel A, Westlund JM, et al. Homotypic clusters of transcription factor binding sites are a key component of human promoters and enhancers. Genome Res. 2010; 20:565–77

6. Lemon B. Orchestrated response: a symphony of transcription factors for gene control. Genes & Development 2000; 14:2551–69.

7. Nolis IK, McKay DJ, Mantouvalou E, et al. Transcription factors mediate long-range enhancer-promoter interactions. Proceedings of the National Academy of Sciences 2009; 106:20222–27.

8. Mendenhall EM, Williamson KE, Reyon D, et al. Locus-specific editing of histone modifications at endogenous enhancers. Nat. Biotechnol. 2013; 31:1133–36.

9. Maurano MT, Haugen E, Sandstrom R, et al. Large-scale identification of sequence variants influencing human transcription factor occupancy in vivo. Nat. Genet. 2015; 47:1393–1401.

10. Jolma A, Taipale J. Methods for analysis of transcription factor DNA-binding specificity in vitro. Subcell Biochem 2011; 52:155—73.

11. Garner MM, Revzin A. A gel electrophoresis method for quantifying the binding of proteins to specific DNA regions: application to components of the Escherichia coli lactose operon regulatory system. Nucleic Acids Res 1981; 9:3047—60.

12. Hampshire AJ, Rusling DA, Broughton-Head VJ, et al. Footprinting: a method for determining the sequence selectivity, affinity and kinetics of DNA-binding ligands. Methods 2007; 42:128—40.

13. Berger MF, Philippakis AA, Qureshi AM, et al. Compact, universal DNA microarrays to comprehensively determine transcription-factor binding site specificities. Nat Biotechnol 2006; 24:1429—35.

14. Jolma A, Kivioja T Toivonen J, et al. Multiplexed massively parallel SELEX for characterization of human transcription factor binding specificities. Genome Res 2010; 20:861—73.

15. Pillai S, Chellappan SP. ChIP on chip and ChIP-seq assays: genome-wide analysis of transcription factor binding and histone modifications. Methods Mol Biol 2015; 1288:447—72.

16. Johnson DS, Mortazavi A, Myers RM, et al. Genome-wide mapping of in vivo protein-DNA interactions. Science 2007; 316:1497—502.

17. Mardis ER. ChIP-seq: welcome to the new frontier. Nat Methods 2007; 4:613—4.

18. Thomas R, Thomas S, Holloway AK, et al. Features that define the best ChIP-seq peak calling algorithms. Brief Bioinform 2017; 18:441—50.

19. Guo Y Mahony S, Gifford DK. High resolution genome wide binding event finding and motif discovery reveals transcription factor spatial binding constraints. PLos Comput Biol 2012; 8:e1002638.

20. Zhang Y, Liu T, Meyer CA, et al. Model-based analysis of ChIP-Seq (MACS). Genome Biol 2008; 9:R137.

21. Worsley Hunt R, Wasserman WW. Non-targetd transcription factors motif are a systemic component of ChIP-seq datasets. Genome Biol 2014; 15:1—16.

22. Pickrell JK, Gaffney DJ, Gilad Y, et al. False positive peaks in ChIP-seq and other sequencing-based functional assays caused by unannotated high copy number regions. Bioinformatics 2011; 27:2144—6.

23. Tognon M, Giugno R, Pinello L. A survey on algorithms to characterize transcription factor binding sites. Brief. Bioinform. 2023; bbad156.

24. Rhee HS, Pugh BF. Comprehensive genome-wide protein-DNA interactions detected at single-nucleotide resolution. Cell 2011; 147:1408—19.

25. John S, Sabo PJ, Thurman RE, et al. Chromatin accessibility pre-determines glucocorticoid receptor binding patterns. Nat Genet 2011; 43(3):264—8.

26. ENCODE Project Consortium. An integrated encyclopedia of DNA elements in the human genome. Nature 2012; 489:57—74.

27. Zheng R, Wan C, Mei S, et al. Cistrome Data Browser: expanded datasetsand new tools for gene regulatory analysis. Nucleic Acids Res 2019; 47(D1):D729—35.

28. Kolmykov S, Yevshin I, Kulyashov M, et al. GTRD: an integrated view of transcription regulation. Nucleic Acids Res 2021; 49(D1):D104—11.

29. Bailey TL, Machanick P. Inferring direct DNA binding from ChIP-seq. Nucleic Acids Res 2012; 40:e128—e128.

30. Worsley Hunt R, Mathelier A, Del Peso L, et al. Improving analysis of transcription factor binding sites within ChIP-Seq data based on topological motif enrichment. BMC genomics 2014; 15:1—19.

31. D’haeseleer P. How does DNA sequence motif discovery work? Nat. Biotechnol. 2006; 24:959–961.

32. Zambelli F, Pesole G, Pavesi G. Motif discovery and transcription factor binding sites before and after the next-generation sequencing era. Brief. Bioinform. 2013; 14:225–237.

33. Stormo GD. DNA binding sites: representation and discovery. Bioinformatics 2000; 16:16–23.

34. Stormo GD. Modeling the specificity of protein-DNA interactions. Quant Biol 2013; 1:115—30.

35. Weirauch MT, Cote A, Norel R, et al. Evaluation of methods for modeling transcription factor sequence specificity. Nat Biotechnol 2013; 31:126–34.

36. Boser BE, Guyon IM, Vapnik VN. A training algorithm for optimal margin classifiers. Proceedings of the fifth annual workshop on Computational learning theory - COLT ’92 1992; 1:144–52.

37. Ben-Hur A, Ong CS, Sonnenburg S, et al. Support Vector Machines and Kernels for Computational Biology. PLoS Computational Biology 2008; 4:e1000173.

38. Boeva V. Analysis of Genomic Sequence Motifs for Deciphering Transcription Factor Binding and Transcriptional Regulation in Eukaryotic Cells. Front. Genet. 2016; 7:24.

39. Lee D, Karchin R, Beer MA. Discriminative prediction of mammalian enhancers from DNA sequence. Genome Res. 2011; 21:2167–80.

40. Ghandi M, Lee D, Mohammad-Noori M, et al. Enhanced Regulatory Sequence Prediction Using Gapped k-mer Features. PLoS Computational Biology 2014; 10:e1003711.

41. Talukder A, Berham C, Li X, et al. Interpretation of deep learning in genomics and epigenomics. Brief Bioinform 2021; 22:bba177.

42. Zeng H, Edwards MD, Liu G, et al. Convolutional neural network architectures for predicting DNA-protein binding. Bioinformatics 2016; 32:i121—7.

43. Alipanahi B, Delong A, Weirauch MT, et al. Predicting the sequence specificities of DNA-and RNA- binding proteins by deep learning. Nat Biotechnol 2015; 33:831—8.

44. Kelley DR, Snoek J, Rinn JL. Basset: learning the regulatory code of the accessible genome with deep convolutional neural networks. Genome Res 2016; 26:990—9.

45. Park S, Koh Y, Jeon H, et al. Enhancing the interpretability of transcription factor binding site prediction using attention mechanism. Sci Rep 2020; 10:13413.

46. Jothi R, Cuddapah S, Barski A, et al. Genome-wide identification of in vivo protein-DNA binding sites from ChIP-Seq data. Nucleic Acids Res 2008; 36:5221—31.

47. Valouev A, Johnson DS, Sundquist A, et al. Genome-wide analysis of transcription factor binding sites based on ChIP-seq data. Nat Methods 2008; 5:829—34.

48. Hu M, Yu J, Taylor JM, et al. On the detection and refinement of transcription factor binding sites using ChIP-Seq data. Nucleic Acids Res 2010; 38:2154—67.

49. Xu J, Gao J, Ni P, et al. Less-is-more: selecting transcription factor binding regions informative for motif inference. Nucleic Acids Res 2024; 52:e20.

50. Shrikumar A, Prakash E, Kundaje A. GkmExplain: fast and accurate interpretation of nonlinear gapped k-mer SVMs. Bioinformatics 2019; 35:i173—i182.

51. Castellana S, Biagini T, Parca L, et al. A comparative benchmark of classic DNA motif discovery tools on synthetic data. Brief in Bioinf 2021; 22:bbab303.

52. Tompa M, Li N, Bailey TL, et al. Assessing computational tools for the discovery of transcription factor binding sites. Nat Biotechnol 2005; 23:137—44.

53. Hombach D, Scwarz JM, Robinson PN, et al. A systematic large-scale comparison of transcription factor binding site models. BMC Genomics 2016; 17:1—10.

54. Ambrosini G, Vorontsov I, Penzar D, et al. Insights gained from a comprehensive all-against-all transcription factor binding motif benchmarking study. Genome Biology 2020; 21:1—18.

55. Bailey TL, Elkan C. Fitting a mixture model by expectation maximization to discover motifs in biopolymers. Proc Int Conf Intell Syst Mol Biol 1994; 2:28–36.

56. Bailey TL, Elkan C. The value of prior knowledge in discovering motifs with MEME. Proc Int Conf Intell Syst Mol Biol 1995; 3:21–9.

57. Bailey TL, Williams N, Misleh C, et al. MEME: discovering and analyzing DNA and protein sequence motifs. Nucleic Acids Res 2006; 34:W369–73

58. Bailey TL. STREME: accurate and versatile sequence motif discovery. Bioinformatics 2021; 37:2834—40.

59. Lee D. LS-GKM: a new gkm-SVM for large-scale datasets. Bioinformatics 2016; 32:2196—98.

60. Mazzocca M, Fillot T, Loffreda A, et al. The needle and the haystack: single molecule tracking to probe the transcription factor search in eukaryotes. Biochemical Society Transactions 2021; 49:1121—32.

61. Stormo GD. Information content and free energy in DNA-protein interactions. Journal of Theoretical Biology 1998; 195:135—7.

62. Schneider TD, Stephens RM. Sequence logos: a new way to display consensus sequences. Nucleic Acids Res 1990; 18:6097—100.

63. Sandelin A, Alkema W, Engström P, et al. JASPAR: an open-access database for eukaryotic transcription factor binding profiles. Nucleic Acids Res 2004; 32:D91—4.

64. Kulakovskiy IV, Medvedeva YA, Schaefer U, et al. HOCOMOCO: a comprehensive collection of human transcription factor binding site models. Nucleic Acids Res 2013; 41:D195—202.

65. Zhao Y, Stormo GD. Quantitative analysis demonstrates most transcription factors require only simple models for specificity. Nat Biotechnol 2011; 29:480—3.

66. Hertz GZ, Stormo GD. Identifying DNA and protein patterns with statistically significant alignments of multiple sequences. Bioinformatics 1999; 15:563—577.

67. Leslie C, eskin E, Noble WS. The spectrum kernel: a string kernel for SVM protein classification. Pac Symp Biocomput 2002; 564—75.

68. Lee D, Kerchin R, Beer MA. Discriminative prediction of mammalian enhancers from DNA sequence. Genome Res 2011; 21:2167—80.

69. Kuang R, Ie E, Wang K, et al. Profile-based string kernels for remote homology detection and motif extraction. Proc IEEE Comput Syst Bioinform Conf 2004; 152—60.

70. Leslie C, Kuang R. Fast Kernels for inexact String Matching. Learning theory and Kernel Machines 2003; 41:W544—56.

71. Agius P, Arvey A, Chang W, et al. High resolution models of transcription factor-DNA affinities improve in vitro and in vivo binding predictions. PLoS Comput Biol 2010; 6:e1000916.

72. Ghandi M, Mohammad-Noori M, Beer MA. Robust k-mer frequency estimation using gapped k-mers. J Math Biol 2014; 69:469—500.

73. Ghandi M, Lee D, Mohammad-Noori M, et al. Enhanced Regulatory Sequence Prediction Using Gapped k-mer Features. PLoS Comput. Biol. 2014; 10:e1003711.

74. Lee DD, Shimko TC, Aditham AK, et al. Comprehensive, high-resolution binding energy landscapes reveal context dependencies of transcription factor binding. Proc Natl Acad Sci USA 2018; 115:E3702—E3711.

75. Meuleman W, Muratov A, Rynes E, et al. Index and biological spectrum of human DNase I hypersensitive sites. Nature 2020; 584:244—51.

76. Song L, Crawford GE. DNase-seq: a high-resolution technique for mapping active gene regulatory elements across the genome from mammalian cells. Cold Spring Harbor Protocols 2010; 2:pdb-prot5384.

77. Krützfeldt LM, Schubach M, Kircher M. The impact of different negative training data on regulatory sequence predictions. PLos One 2020; 15:e0237412.

78. Bailey TL. DREME: motif discovery in transcription factor ChIP-seq data. Bioinformatics 2011; 27:1653—9.

79. Heinz S, Benner C, Spann N, et al. Simple combinations of lineage-determining transcription factors prime cis-regulatory elements required for macrophage and B cell identities. Mol Cell 2010; 38:576—89.

80. Grant CE, Bailey TL, Noble WS. Fimo: scanning for occurrences of a given motif. Bioinformatics 2011; 27:1017—18.

81. Jalili V, Matteucci M, Masseroli M, et al. Using combined evidence from replicates to evaluate ChIP-seq peaks. Bioinformatics 2015; 31:2761—69.

82. Sand O, Turatsinze JV, van Helden J. Evaluating the prediction of cis-acting regulatory elements in genome sequences. Modern Genome Annotation: The BioSapiens Network 2008; 55—89.

83. Samee MAH, Bruneau BG, Pollard KS. A de novo shape motif discovery algorithm reveals preferences of transcription factors for DNA shape beyond sequence motifs. Cell systems 2019; 8:27—42

84. Jayaram N, Usvyat D, R. Martin AC. Evaluating tools for transcription factor binding site prediction. BMC bioinf 2016; 17:1—12.

85. Bell AC, West AG, Felsenfeld G. The protein CTCF is required for the enhancer blocking activity of vertebrate insulators. Cell 1999; 98:387—96.

86. Tessari MA, Gostissa M, Altamura S, et al. Transcriptional activation of the cyclin A gene by the architectural transcription factor HMGA2. Molecular and cellular biology 2003; 23:9104—16.

87. Lazaro-Camp VJ, Salari K, Meng X, et al. SETDB1 in cancer: overexpression and its therapeutic implications. American journal of cancer research 2021; 11:1803.

88. Musa J, Aynaud MM, Mirabeau O, et al. MYBL2 (B-Myb): a central regulator of cell proliferation, cell survival and differentiation involved in tumorigenesis. Cell death & disease 2017; 8:e2895—e2895.

89. Lefebvre V. The SoxD transcription factors—Sox5, Sox6, and Sox13—are key cell fate modulators. The international journal of biochemistry & cell biology 2010; 42:429—32.

90. Yagi R, Kohn MJ, Karavanova I, et al. Transcription factor TEAD4 specifies the trophectoderm lineage at the beginning of mammalian development. Development 2007; 134:3827—36.

91. Stringham JL, Brown AS, Drewell RA, et al. Flanking sequence context-dependent transcription factor binding in early Drosophila development. BMC bioinformatics 2013; 14:1—13.

92. Boytsov A, Abramov S, Makeev VJ, et al. Positional weight matrices have sufficient prediction power for analysis of noncoding variants. F1000Research 2022; 11.

93. Üstün B, Melssen WJ, Buydens LMC. Visualization and interpretation of support vector regression models. Analytics chimica acta 2007. 595:299—309.

94. Guido R, Ferrisi S, Lofaro D, et al. An overview on the advancements of support vector machine models in healthcare applications: a review. Information 2024; 15:235.

95. Srivastava D, Mahony S. Sequence and chromatin determinants of transcription factor binding and the establishment of cell-type specific binding patterns. Biochimica et Biophysica Acta (BBA)-Gene Regulatory Mechanisms 2020; 1863:194443.

96. Awdeh A, Turcotte M, Perkins TJ. Identifying transcription factor with cell-type specific DNA binding signatures. BMC Genomics 2024; 25:957.

97. Gertz J, Savic D, Varley KE, et al. Distinct properties of cell-type specific and shared transcription factor binding sites. Molecular Cell 2013; 52:25—36.

98. Sun N, Zhao H. Reconstructing transcriptional regulatory networks through genomics data. Statistical methods in medical research 2009; 18:595—617.

99. Baur B, Shin J, Zhang S, et al. Data integration for inferring context-specific gene regulatory networks. Current opinion in systems biology 2020; 23:38—46.

100. Park PJ. ChIP-seq: advantages and challenges of a maturing technology. Nature reviews genetics 2009; 10:669—80.

101. Awdeh A, Turcotte M, Perkins TJ. WACS: improving ChIP-seq peak calling by optimally weighting controls. BMC bioinformatics 2021; 22:1—21.

102. Furey TS. ChIP-seq and beyond: new and improved methodologies to detect and characterize protein-DNA interactions. Nature reviews genetics 2012; 13:840—52.

103. Levo M, Zalckvar E, Sharon E, et al. Unraveling determinants of transcription factor binding outside the core binding site. Genome research 2015; 25:1018—29.

104. Slattery M, Zhou T, Yang L, et al. Absence of a simple code: how transcription factors read the genome. Trends in biochemical sciences 2014; 39:381—99.

## References

1. Bailey TL. STREME: accurate and versatile sequence motif discovery. Bioinformatics 2021; 37:2834—40.

2. Zambelli F, Pesole G, Pavesi G. Motif discovery and transcription factor binding sites before and after the next-generation sequencing era. Brief Bioinform 2013; 14:225–37.

3. Tognon M, Giugno R, Pinello L. A survey on algorithms to characterize transcription factor binding sites. Brief Bioinform 2023; bbad156

4. Stormo GD. DNA binding sites: representation and discovery. Bioinformatics 2000; 16:16–23.

5. Stormo GD. Modeling the specificity of protein-DNA interactions. Quant Biol 2013; 1:115—30.

6 Weiner P. Linear Pattern Matching Algorithms. 14^th^ Annual Symposium on Switching and Automata Theory 1973; 1:1–11.

7. Bailey TL, Elkan C. Fitting a mixture model by expectation maximization to discover motifs in biopolymers. Proc Int Conf Intell Syst Mol Biol 1994; 2:28–36.

8. Bailey TL, Elkan C. The value of prior knowledge in discovering motifs with MEME. Proc Int Conf Intell Syst Mol Biol 1995; 3:21–9.

9. Bailey TL, Williams N, Misleh C, et al. MEME: discovering and analyzing DNA and protein sequence motifs. Nucleic Acids Res 2006; 34:W369–73

10. Hashim FA, Mabrouk MS, Al-Atabany W. Review of different sequence motif finding algorithms. Avicenna journal of medical biotechnology 2019; 11:130.

11. Lee D. LS-GKM: a new gkm-SVM for large-scale datasets. Bioinformatics 2016; 32:2196—98.

12. Ghandi M, Mohammad-Noori M, Beer MA. Robust k-mer frequency estimation using gapped k-mers. J Math Biol 2014; 69:469—500.

13. Ghandi M, Lee D, Mohammad-Noori M, Beer MA. 2014. Enhanced Regulatory Sequence Prediction Using Gapped k-mer Features. PLoS Comput Biol 10: e1003711.

14. Shrikumar A, Prakash E, Kundaje A. GkmExplain: fast and accurate interpretation of nonlinear gapped k-mer SVMs. Bioinformatics 2019; 35:i173—i182.

15. ENCODE Project Consortium. An integrated encyclopedia of DNA elements in the human genome. Nature 2012; 489:57—74.

16. Lambert SA, Jolma A, Campitelli LF, et al. The Human Transcription Factors. Cell 2018; 172:650–65.

17. Quinlan AR, Hall IM. BEDTools: a flexible suite of utilities for comparing genomic features. Bioinformatics 2010; 26:841—42.

18. Meuleman W, Muratov A, Rynes E, et al. Index and biological spectrum of human DNase I hypersensitive sites. Nature 2020; 584:244—51.

19. Bailey TL, Bodén M, Buske FA, et al. MEME SUITE: tools for motif discovery and searching, Nuc Acids Res 2009;37:W202—8.

20. Jiang M, Anderson J, Gillespie J, et al. uShuffle: A useful tool for shuffling biological sequences while preserving the k-let counts. BMC Bioinformatics 2008;9:192.

21. Propp JG, Wilson DB. How to get a perfectly random sample from a generic Markov chain and generate a random spanning tree of a directed graph. Journal of Algorithms 1998;27:170–217.

22. Wilson DB. Generating random spanning trees more quickly than the cover time. Proceedings of the 28th Annual ACM Symposium on the Theory of Computing (STOC’96) 1996; 296–303.

23. Grant CE, Bailey TL, Noble WS. Fimo: scanning for occurrences of a given motif. Bioinformatics 2011; 27:1017—18.

24. Fedotova AA, Bonchuk AN, Mogila VA, et al. C2H2 zinc finger proteins: the largest but poorly explored family of higher eukaryotic transcription factors. Acta Naturae 2017; 9:47—58.

25. Bell AC, West AG, Felsenfeld G. The protein CTCF is required for the enhancer blocking activity of vertebrate insulators. Cell 1999; 98:387—96.

26. Ferreira R, Ohneda K, Yamamoto M, et al. GATA1 function, a paradigm for transcription factors in hematopoiesis. Molecular and cellular biology 2005.

27. Vicente C, Conchillo A, García-Sánchez MA, et al. The role of the GATA2 transcription factor in normal and malignant hematopoiesis. Critical reviews in oncology/hematology 2012; 82:1—17.

28. Smart DE, Green K, Oakley F, et al. JunD is a profibrogenic transcription factor regulated by Jun N-terminal kinase-independent phosphorylation. Hepatology 2006; 44:1432—1440.

29. Tijssen, Marloes R et al. Genome-wide analysis of simultaneous GATA1/2, RUNX1, FLI1, and SCL binding in megakaryocytes identifies hematopoietic regulators. Developmental cell vol. 20,5 (2011): 597-609.

30. Srivastava, P.K., Hull, R.P., Behmoaras, J. et al. JunD/AP1 regulatory network analysis during macrophage activation in a rat model of crescentic glomerulonephritis. BMC Syst Biol 7, 93 (2013).

31. Fischer M, Steiner L, Engeland K. The transcription factor p53: not a repressor, solely an activator. Cell cycle 2014; 13:3037—58.

